# α-Actinin-1 promotes activity of the L-type Ca^2+^ Channel Ca_V_1.2

**DOI:** 10.1101/664102

**Authors:** Matthew Turner, David E. Anderson, Madeline Nieves-Cintron, Peter Bartels, Andrea M. Coleman, Peter B. Henderson, Kwun Nok Mimi Man, Vladimir Yarov-Yarovoy, Donald M. Bers, Manuel F. Navedo, Mary C. Horne, James B. Ames, Johannes W. Hell

## Abstract

The L-type Ca^2+^ channel Ca_V_1.2 governs gene expression, cardiac contraction, and neuronal activity. Binding of α-actinin to the IQ motif of Ca_V_1.2 supports its surface localization and postsynaptic targeting in neurons. We report a bi-functional mechanism that restricts Ca_V_1.2 activity to its target sites. We solved separate NMR structures of the IQ motif (residues 1646-1664) bound to α-actinin-1 and to apo-calmodulin (apoCaM). The Ca_V_1.2 K1647A and Y1649A mutations, which impair α-actinin-1 but not apoCaM binding, but not the F1658A and K1662E mutations, which impair apoCaM but not α-actinin-1 binding, decreased single channel open probability, gating charge movement, and its coupling to channel opening. Thus, α-actinin recruits Ca_V_1.2 to defined surface regions and simultaneously boosts its open probability so that Ca_V_1.2 is mostly active when appropriately localized.

## INTRODUCTION

Ca^2+^ influx through Ca_V_1.2 is critical for the functions of many organs as strikingly illustrated by Timothy syndrome (Splawski et al., 2004). In this disease a point mutation in Ca_V_1.2 causes, among other symptoms, lethal arrhythmias, autistic-like behaviors, immune deficiency, and webbing of fingers (Splawski et al., 2004). Ca_V_1.2 is the main L-type channel in heart (Seisenberger et al., 2000), vascular smooth muscle cells (Ghosh et al., 2017), and brain (Hell et al., 1993, Sinnegger-Brauns et al., 2004). Ca^2+^ influx through Ca_V_1.2 triggers cardiac contraction, regulates arterial tone (Ghosh et al., 2017), mediates different forms of synaptic long-term potentiation (Grover and Teyler, 1990, Patriarchi et al., 2016, Qian et al., 2017), and controls neuronal excitability (Berkefeld et al., 2006, Marrion and Tavalin, 1998). Furthermore, L-type channels are much more strongly coupled to gene expression than other Ca^2+^ channels (Cohen et al., 2018, Dolmetsch et al., 2001, Li et al., 2012, Ma et al., 2014). Finally, Ca_V_1.2 forms a physical and functional complex with the β_2_ adrenergic receptor (Davare et al., 2001) making it a prime target for signaling by norepinephrine (Patriarchi et al., 2016, Qian et al., 2017), which is important for wakefulness, attention, and various forms of learning (Berman and Dudai, 2001, Cahill et al., 1994, Carter et al., 2010, Hu et al., 2007, Minzenberg et al., 2008).

Ca_V_1.2 consists of the pore-forming α_1_1.2 subunit and auxiliary α_2_δ and β subunits, which facilitate release from the endoplasmic reticulum and the controlled trafficking of Ca_V_1.2 to the cell surface (Dai et al., 2009, Dolphin, 2012, 2016, Ghosh et al., 2018, Zamponi et al., 2015). However, α_2_δ and β subunits do not target Ca_V_1.2 to specific sites in the plasma membrane. Rather, Ca_V_1.2 anchoring at defined regions at the cell surface is mediated by α-actinin, which binds to the IQ motif in the C-terminus of α_1_1.2 (Hall et al., 2013, Tseng et al., 2017).

A systematic yeast two hybrid screen defined three residues in the IQ motif of α_1_1.2, whose mutations to alanine residues affect α-actinin binding: K1647A, Y1649A, and I1654A. All three mutations reduced surface expression of Ca_V_1.2 by ∼35% but current density by 70-80% (Tseng et al., 2017). These results suggest that α-actinin binding to the IQ motif promotes not only surface localization but also channel activity. Such a multifunctional role would ensure that Ca_V_1.2 is mostly active at its ultimate destinations and much less so when in transit and outside its target areas.

The closely related α_1_1.3 subunit of the L-type channel Ca_V_1.3 shares nearly 100 % sequence identity with α_1_1.2 in its membrane proximal 165 residues of its C-terminus in which the IQ motif is embedded (the eponymous Ile is I1654 of α_1_1.2 and I1609 of α_1_1.3; Suppl. Fig. 1). Ca^2+^-free calmodulin (apoCaM) binds to this IQ motif and mutation of I1609 in α_1_1.3 impairs both, apoCaM binding and open probability Po of Ca_V_1.3 (Adams et al., 2014, Ben Johny et al., 2013, Ben-Johny et al., 2014). We solved the NMR structures of the third and fourth EF-hands of α-actinin-1 (EF3 and EF4, residues 822-892; Fig. 1A) and of full length apoCaM bound to the α-helical IQ motif of Ca_V_1.2 (residues 1646-1664). This work provided new insight into the structure of Ca_V_1.2 especially as relevant for these two critical binding partners and informed experiments that dissected the exact functions of α-actinin versus apoCaM binding. Refined analysis of Ca_V_1.2 activity by cell attached single channel recording revealed that point mutations that affected α-actinin-1 but not those that affected apoCaM binding dramatically decreased the channel Po by impairing gating charge movement as well as its coupling to channel opening. We conclude that α-actinin plays a dual role by anchoring Ca_V_1.2 at specific subcellular domains such as the postsynaptic sites and at the same time boosting its open probability. This mechanism ensures that activity of Ca_V_1.2 is minimal when in transit during secretory trafficking and outside its intended location at the cell surface, where its Ca^2+^ conductance could adversely affect cell functions, but maximal at its final destination.

**Fig. 1.**
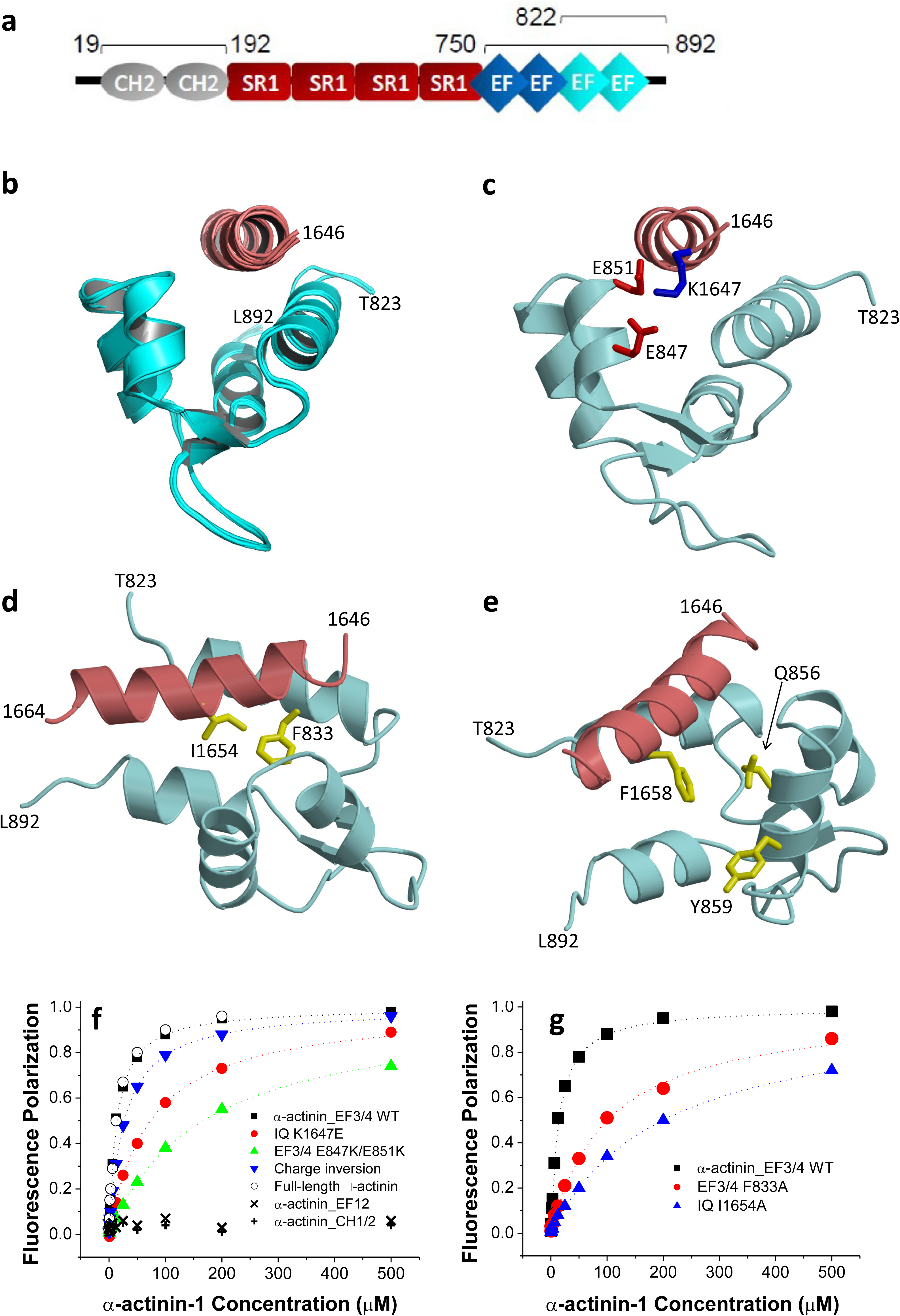
α-Actinin-1 makes electrostatic and hydrophobic contacts with Ca_V_1.2. a, Linear model of α-actinin domains (CH, calponin homology domains; SR, spectrin repeats; EF, EF hands). Bars and numbers on top of the model depict the segments used in this work. b, Ensemble of 10 lowest energy NMR-derived structures of α-actinin-1_EF34 (cyan) bound to IQ peptide (red). Structural statistics are given in Table 1. c, Energy minimized average structure of α-actinin-1_EF34 (cyan) bound to IQ (red), revealing intermolecular salt bridges between K1647 of IQ and E847/E851 of α-actinin-1. The K1647 side chain amino nitrogen atom is 2.8 Å and 2.5 Å away from the side chain carbonyl oxygen atoms of E847 and E851, respectively. d, Intermolecular hydrophobic contacts between I1654 of IQ (red) and F833 of α-actinin-1 (cyan). The I1654 side chain methyl carbon atom is 2.9 Å away from the closest aromatic ring atom of F833. Side-chain atoms are colored yellow. e, Lack of direct contacts between F1658 of IQ (red) and α-actinin-1 (cyan). The aromatic side chain of F1658 is primarily solvent exposed and does not directly contact α-actinin-1. The aromatic ring of F1658 is closest to the aromatic ring of Y859 and β-methylene carbon of Q856 of α-actinin-1, which are 5.2 Å and 4.3 Å apart, respectively. f, FP titrations show binding of WT α-actinin-1_EF34 to IQ peptides WT (black) and K1647E (red), of α-actinin-1_EF34 mutant E847K/E851K to IQ peptides WT (green) and K1647E (purple), of full-length α-actinin-1 to IQ WT (o), and lack of IQ binding to α-actinin-1_CH1-CH2 (**+**) and α-actinin-1_EF12 (x; see Table 2 for binding parameters and standard errors). g, FP titrations showing binding of wild type α-actinin-1_EF34 to IQ peptides WT (black) and I1654A (purple) and of ACTN1_EF34 mutant F833 to IQ WT (red; see Table 2 for binding parameters and standard errors).

**Table 1:** NMR Structural Statistics for α-actinin-1 EF34/IQ

| **NMR Structural Restraints** |  |
| --- | --- |
| Intermolecular NOEs | 73 |
| Hydrogen Bonds | 62 |
| RDC Q-Factor | 0.095 |
| $^1D_{HN}RDC$ | 34 |
| RDC Correlation Coefficient (R) | 0.99 |
| **Root mean squared deviation from average structure** |  |
| α-actinin-1 backbone atoms | $0.5 \text{ \AA} \pm 0.3$ for 200 structures |
| α-actinin-1 backbone atoms (refined) | $0.3 \text{ \AA} \pm 0.2$ for 50 structures |
| **Haddock Scoring** |  |
| Cluster Size | 194 |
| Van der Waals Energy | $-40.2 \pm 6.2$ |
| Electrostatic Energy | $-395.4 \pm 37.1$ |
| Restraints Violation Energy | $278.0 \pm 6.23$ |
| **Ramachandran Plot** |  |
| Most Favored Regions | 96.7 % |
| Allowed Regions | 2.2 % |
| Unfavored Regions | 1.1 % |

**Table 2:** Kd (in μM) for α-actinin-1 EF3/EF4 binding to IQ as determined by FP. Shown are mean+SD (n=3 for all conditions)

| | | α-actinin-1 EF3/4 | | | |
| --- | --- | --- | --- | --- | --- |
|  |  | WT | E847K/E851K | F833A |  |
| Cav1.2 IQ | WT | 14 ± 2 | 167 ± 20 | 100 ± 10 | |
| | K1647E | 73 ± 10 | 27 ± 2 | / | |
| | K1647A | 67 ± 10 | / | / | |
| | Y1649A | 42 ± 4 | / | / | |
| | I1654A | 200 ± 20 | / | / | |
| | F1658A | 20 ± 2 | / | / | |
| | K1662E | 20 ± 2 | 150 ± 20 | / | |

## RESULTS

### Binding of α-actinin-1 EF3/EF4 to the Ca_V_1.2 IQ motif

α-actinin is encoded by four homologous genes with α-actinin-1 and α-actinin-2 being most prominent in neurons (Hall et al., 2013, Hell, 2014, Matt et al., 2018, Wyszynski et al., 1997). α-actinins consist of two N-terminal calponin homology domains (CH1, CH2; residues 19-192), four central spectrin repeats (SR1-4), which form a rod-like coiled-coil structure (Ribeiro Ede et al., 2014), and four EF hands at their C-termini (residues 750-892; Fig. 1a). We first performed NMR spectroscopy with the CH1/CH2 region (residues 19-192). Two-dimensional NMR spectra of ^15^N-labeled CH1/CH2 recorded in the presence and absence of unlabeled IQ peptide appeared to be virtually identical, consistent with a lack of IQ binding to CH1/CH2 under NMR conditions (Suppl. Fig. 2a). The lack of IQ binding to the CH1/CH2 domain is supported by absence of detectable binding measured by fluorescence polarization (see below Fig. 1f). In stark contrast, the NMR spectrum of ^15^N-labeled EF-hand domain of α-actinin-1 (residues 750-892) showed clearly detectable spectral changes upon adding a saturating amount of unlabeled IQ peptide, demonstrating a binding interaction (Suppl. Fig. 2b). The NMR peaks of α-actinin-1 most affected by the binding of IQ (see labeled peaks in Suppl. Fig. 2b) were assigned to residues in EF3 and EF4 (the C-lobe of the EF hand region; residues 822-892). Consistently, the NMR spectrum of ^15^N-labeled α-actinin-1 EF3/4 (residues 822-892) exhibited large spectral changes upon addition of the IQ peptide (Suppl. Fig. 2c), which are similar to those seen with the construct that contains all four EF-hands (Suppl. Fig. 2b). We conclude that the IQ peptide binds to the α-actinin-1 C lobe.

### NMR Structure of α-actinin-1 EF3/4 bound to the IQ motif of Ca_V_1.2

We had previously reported NMR spectral assignments for α-actinin-1 EF3/4 (BMRB accession number 25902) (Turner et al., 2016). We used these assignments to obtain NMR-derived structural restraints for high-resolution structural analysis of α-actinin-1 EF3/4 bound to the Ca_V_1.2 IQ motif (EF3/4-IQ). Atomic-level NMR structures were calculated on the basis of distance restraints derived from analysis of NOESY spectra and long-range orientational restraints derived from residual dipolar coupling (RDC) data (Suppl. Fig. 3a). EF34 in the complex was resolved for 69 residues, starting at T823 and ending at L892. The 10 lowest-energy NMR structures are overlaid in Fig. 1b and structural statistics summarized in Table 1. The overall precision of the ensemble was expressed by an RMSD of 0.3 Å calculated from the coordinates of the main chain atoms. The energy-minimized average structure of EF3/4-IQ (Fig. 1c, calculated from the ensembles in Fig. 1b) contained two EF-hand motifs (α1: A826 – A837; α2: M845 – E851; α3: P854 – R863; α4: M880 – Y887; β1: Y842 – T844; β2: A876 – D878) bound to an α-helical IQ motif (Ca_V_1.2 residues 1646-1664). The NMR structure of EF3/4-IQ (Fig. 1c) was quite similar (1.8 Å RMSD) to the NMR structure of α-actinin-2 bound to the seventh Z-repeat of titin (Atkinson et al., 2001). It contained important intermolecular contacts that stabilize the EF3/4 – IQ interaction (Fig. 1c-e). Most striking were a salt bridge between IQ K1647 and EF3/4 E847/E851 (Fig. 1c) and hydrophobic contacts involving IQ I1654 and EF3 F833 (Fig. 1d). The IQ residue F1658 was mostly solvent exposed in the EF3/4-IQ complex contributing minimally if at all to the EF3/4 – IQ interaction (Fig. 1e), in contrast to being buried inside apoCaM in the apoCaM/IQ complex (see below).

### Validation of the α-actinin-1 EF3/4 - IQ NMR structure by mutagenesis

Mutations in both α-actinin-1 EF3/4 and IQ peptide were designed to verify their predicted intermolecular contacts. A synthetic fluorescein-labeled IQ peptide (residues 1644-1666) was titrated with EF3/4 and fluorescence polarization (FP) monitored to determine their K_d_ values. The IQ peptide bound to α-actinin-1 EF3/4 with nearly the same affinity as to full-length α-actinin-1 (Fig. 1f; Table 2). The IQ peptide did not bind to α-actinin-1 EF1/2 (Fig. 1f). Accordingly, IQ interacts exclusively with the C-lobe but not N-lobe of the EF hand domain in α-actinin-1.

The salt bridges formed between IQ residue K1647 and α-actinin-1 residues E847 and E851 (Fig. 1c) provided an opportunity for charge inversion experiments. The single residue alterations K1647A and K1647E in the IQ peptide increased the K_d_ by ∼5-fold and mutating E847 and E851 in α-actinin-1 EF3/4 to lysine (K) by ∼12-fold (Fig. 1f; Table 2). Combining the E847K/E851K mutations in α-actinin-1 EF3/4 with the K1647E substitution in the IQ peptide mostly but not fully restored the binding affinity to the K1647E IQ peptide. This powerful charge inversion experiment unequivocally identified the importance of those salt bridges for the α-actinin-1 – IQ interaction. Furthermore, I1654 of the IQ motif was predicted to form hydrophobic interactions with F833 in EF3 (Fig. 1d). In fact, the α-actinin-1 EF3/4 mutant F833A showed an ∼8-fold decrease in binding affinity, which is consistent with the decrease observed for the I1654A substitution in the IQ peptide (Fig. 1g; Table 2). These results confirmed two important intermolecular interactions that had been seen in the NMR structure (Fig. 1c): a salt bridge contact between K1647 (Ca_V_1.2) and E847/E851 (α-actinin-1) as well as the linchpin hydrophobic contact between I1654 (Ca_V_1.2) and F833 (α-actinin-1).

### NMR Structure of apoCaM bound to the IQ motif of Ca_V_1.2

ApoCaM is predicted to bind to the IQ motif of Ca_V_1.3 under basal conditions to augment Po (Adams et al., 2014) but no structure has so far been available to aid data interpretation. The ^1^H-^15^N HSQC NMR spectrum of ^15^N-labeled apoCaM showed detectable spectral changes upon adding a saturating amount of unlabeled IQ peptide, demonstrating binding (Suppl. Fig. 2d). The NMR peaks of apoCaM most affected by the binding of IQ (see labeled peaks in Suppl. Fig. 2d) were assigned to residues in its C lobe (CaM EF3/4). Complete NMR spectral assignments for apoCaM bound to the IQ peptide had been reported previously (Lian et al., 2007). We used these assignments to obtain NMR-derived structural restraints and determine the atomic-level NMR structure of apoCaM bound to the IQ peptide (called apoCaM-IQ). NMR-derived structures were calculated on the basis of NOESY distance restraints and long-range orientational restraints derived from NMR RDC data (Suppl. Fig. 3b) (Tjandra and Bax, 1997). Our NMR chemical shift analysis indicated that the IQ peptide contacted residues in the C-lobe (residues 82-148) but not N-lobe of apoCaM (residues 1-78; Fig. 2). The 10 lowest-energy NMR structures are overlaid in Fig. 2a and structural statistics summarized in Table 3. The overall precision of the ensemble was expressed by an RMSD of 0.6 Å calculated from the coordinates of the main chain atoms. The energy-minimized average structure of apo-CaM/IQ (Fig. 2b, calculated from the ensembles in Fig. 2a) contained two EF-hand motifs (EF3 and EF4) in a semi-open conformation akin to that observed for apoCaM bound to the IQ motif in voltage-gated Na^+^ channels (Chagot and Chazin, 2011, Feldkamp et al., 2011). This semi-open apoCaM C-lobe structure was bound to the α-helical IQ motif (α_1_1.2 residues 1646-1665; Fig. 2b) with an orientation that was opposite to that observed in the crystal structure of Ca^2+^-CaM bound to IQ (Van Petegem et al., 2005).

**Fig. 2.**
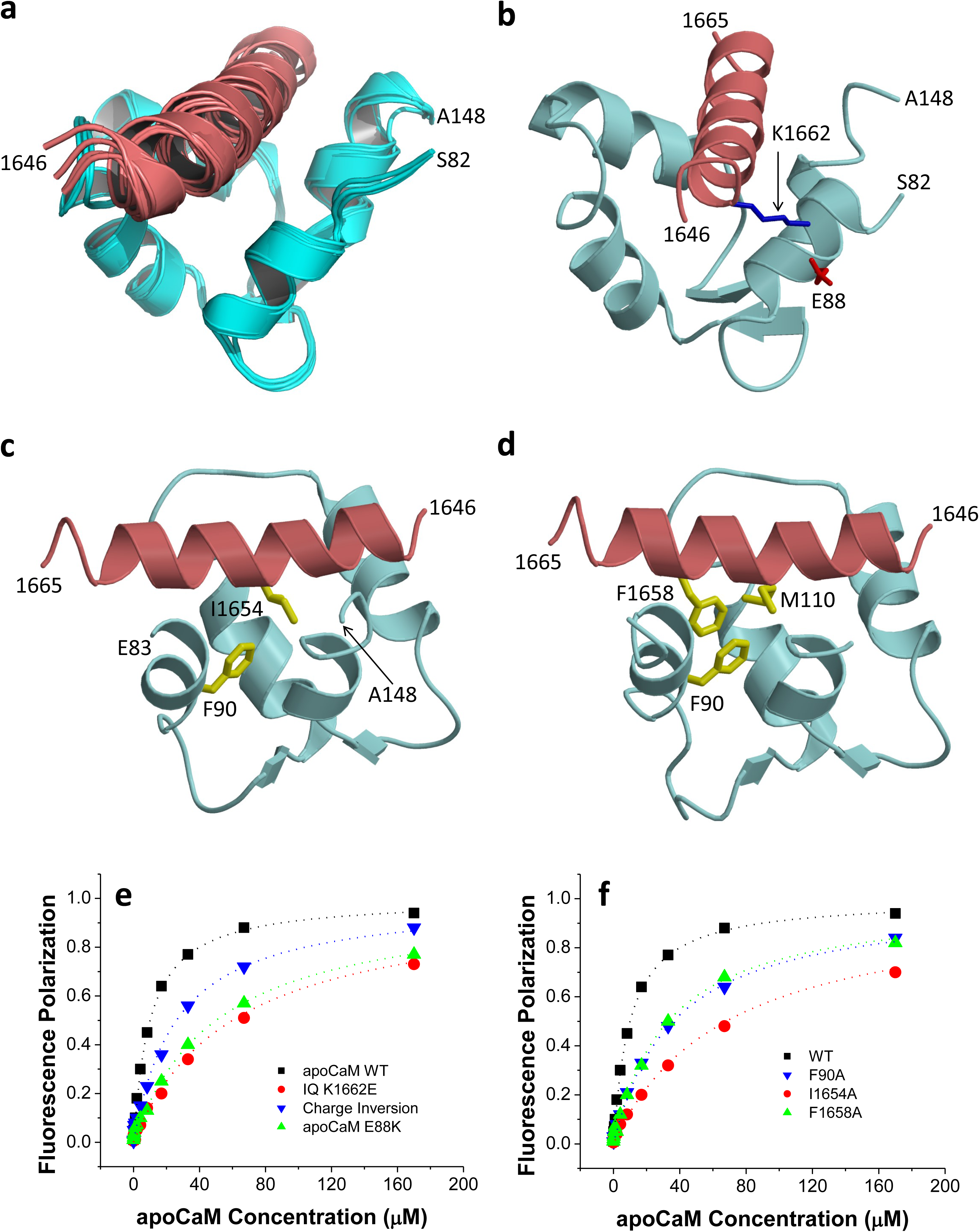
ApoCaM makes electrostatic and hydrophobic contacts with Ca_V_1.2. a, Ensemble of 10 lowest energy NMR-derived structures of apo CaM/IQ complex. Structural statistics are given in Table 3. b, Energy minimized average structure of apoCaM (cyan) bound to IQ (red), showing an intermolecular salt bridge between IQ K1662 and apoCaM E88. The K1662 side chain amino nitrogen atom is 2.7 Å away from the side chain carbonyl oxygen of E88. c, Intermolecular hydrophobic contacts between IQ I1654 (red) and apoCaM F90 (cyan). Side chain atoms colored yellow. The I1654 side chain methyl carbon atom is 2.2 Å away from the closest aromatic ring atom of F90. d, Intermolecular hydrophobic contacts between F1658 of IQ and F90/M110 in apoCaM. The aromatic side chain atoms of F1658 are 2.3Å and 2.6 Å away from the closest side chain atom of F90 and M110, respectively. e, FP titrations showing binding of apoCaM to IQ peptides WT (black) and K1662E (red) and of apoCaM mutant E88K to IQ WT (green) and IQ peptide K1662E (purple, “charge inversion”; see Table 4 for binding parameters and standard errors). f, FP titrations showing binding of apo-CaM to IQ peptides WT (black), I1654A (red), and F1658A (green) and of apoCaM mutant F90A to IQ WT (blue; see Table 4 for binding parameters and standard errors).

**Table 3:** NMR Structural Statistics for apo CaM/IQ

| **NMR Structural Restraints** |  |
| --- | --- |
| Hydrogen Bonds | 62 |
| RDC Q-Factor | 0.089 |
| $^1D_{HN}RDC$ | 36 |
| RDC Correlation Coefficient (R) | 0.99 |
| **Root mean squared deviation from average structure** |  |
| CaM backbone atoms | $0.6 \text{ \AA} \pm 0.3$ for 200 structures |
| CaM backbone atoms (refined) | $0.6 \text{ \AA} \pm 0.2$ for 92 structures |
| **Haddock Scoring** |  |
| Cluster Size | 200 |
| Van der Waals Energy | $-56.8 \pm 0.6$ |
| Electrostatic Energy | $-463.5 \pm 20.7$ |
| Restraints Violation Energy | $99.8 \pm 36.2$ |
| **Ramachandran Plot** |  |
| Most Favored Regions | 89.8 % |
| Allowed Regions | 8.0 % |
| Unfavored Regions | 2.3 % |

**Table 4:** Kd (in μM; means±SEM) for apoCaM binding to IQ as determined by FP. Shown are mean+SD (n=3 for all conditions)

|  |  | apoCaM |  |  |  |
| --- | --- | --- | --- | --- | --- |
|  |  | WT | E88K | F90A |  |
| Cav1.2 IQ | WT | 10 ± 3 | 50 ± 10 | 33 ± 5 | |
| | K1647E | 12 ± 3 | 53 ± 10 | / | |
| | K1647A | 11 ± 3 | / | / | |
| | Y1649A | 16 ± 4 | / | / | |
| | I1654A | 68 ± 10 | / | / | |
| | F1658A | 33 ± 5 | / | / | |
| | K1662E | 60 ± 10 | 26 ± 5 | / | |

The NMR-derived structure of apoCaM bound to α_1_1.2 IQ peptide contained a number of important intermolecular contacts that stabilized this interaction (Fig. 2b-d). The most striking intermolecular interactions were: (1) a salt bridge between IQ residue K1662 and apoCaM residue E88 (Fig. 2b); (2) hydrophobic contacts involving the side chain atoms of IQ residue I1654 and apoCaM residue F90 (Fig. 2c); and (3) hydrophobic contacts involving the side chain atoms of IQ residue F1658 and apoCaM residues F90/M110 (Fig. 2d).

### Validation of the apoCaM - IQ NMR structure by mutagenesis

Mutations in both apoCaM and IQ peptide were introduced to verify their predicted intermolecular contacts. We titrated fluorescein-labeled IQ peptide (residues 1644-1666) with apoCaM and measured FP. The Kd for WT IQ peptide was 10 μM (Fig. 2e, Table 4), similar to the Kd of 13 μM in earlier work (Evans et al., 2011). To understand why this Kd is ∼20-fold higher than the Kd of 580 nM deducted previously from isothermal titration calorimetry (ITC) measurements (Findeisen et al., 2013), we performed ITC experiments by adding increments of apoCaM (100 μM) to the IQ peptide (10 μM) under near physiological salt concentration (100 mM KCl, Suppl. Fig 4a). No heat signal other than that of dilution was detectable (Suppl. Fig. 4b). The prior ITC experiments were performed at 5 mM KCl (Findeisen et al., 2013) and may represent a non-physiological electrostatic attraction between oppositely charged apoCaM and IQ that is suppressed by more physiological salt levels (100 mM KCl). Indeed, apoCaM binds to the IQ peptide with nearly 4-fold higher affinity in the absence of salt (Suppl. Fig. 4c; Kd is 2.6 μM at 0 KCl and 10 μM at 100 mM KCl).

According to our FP binding assay, the K1662E mutation in the IQ peptide and the E88K mutation in apoCaM each decreased the binding affinity between IQ peptide apoCaM by more than 5-fold (Fig. 2e; Table 4). Combining the apoCaM E88K mutation with the IQ peptide alteration K1662E restored the binding affinity to some degree although not completely (Fig. 2e). The apoCaM mutant F90A showed an ∼3.3-fold decrease in binding affinity for WT IQ peptide (Fig. 2f; Table 4). Consistently, I1654A and F1658A substitutions in the IQ peptide resulted in a comparable reduction in their affinity for apoCaM (Fig. 2f). These results confirmed three important intermolecular interactions seen in the NMR structure: (1) a salt bridge between IQ residue K1662 and apoCaM residue E88; (2) hydrophobic contact between IQ residue I1654 and apoCaM residue F90; and (3) hydrophobic contact between IQ residue F1658 and apoCaM residues F90 and M110.

### α-Actinin binding to the Ca_V_1.2 IQ motif augments open probability of individual channels

Point mutations that impaired α-actinin binding to the IQ motif of α_1_1.2 reduced current density upon reconstitution of Ca_V_1.2 in HEK293 cells by 70-80% but surface expression by only 35-40% as determined by two different methods (Tseng et al., 2017). We performed cell attached recordings to precisely determine single channel parameters of Ca_V_1.2 as before (Bartels et al., 2018, Davare et al., 2001, Patriarchi et al., 2016, Qian et al., 2017). All three mutations in the IQ motif that impaired α-actinin binding (Table 2) (Tseng et al., 2017), i.e., K1647A, Y1649A, and I1654A, decreased functional availability by 79-93% and overall single channel activity by ∼85-92% (Fig. 3a-c; Suppl. Fig. 5, 6a,b; Table 5). Ensemble averages of each experiments were similarly decreased by 84-91% (Fig. 3b,e, Suppl. Fig. 5,6d).

**Fig. 3.**
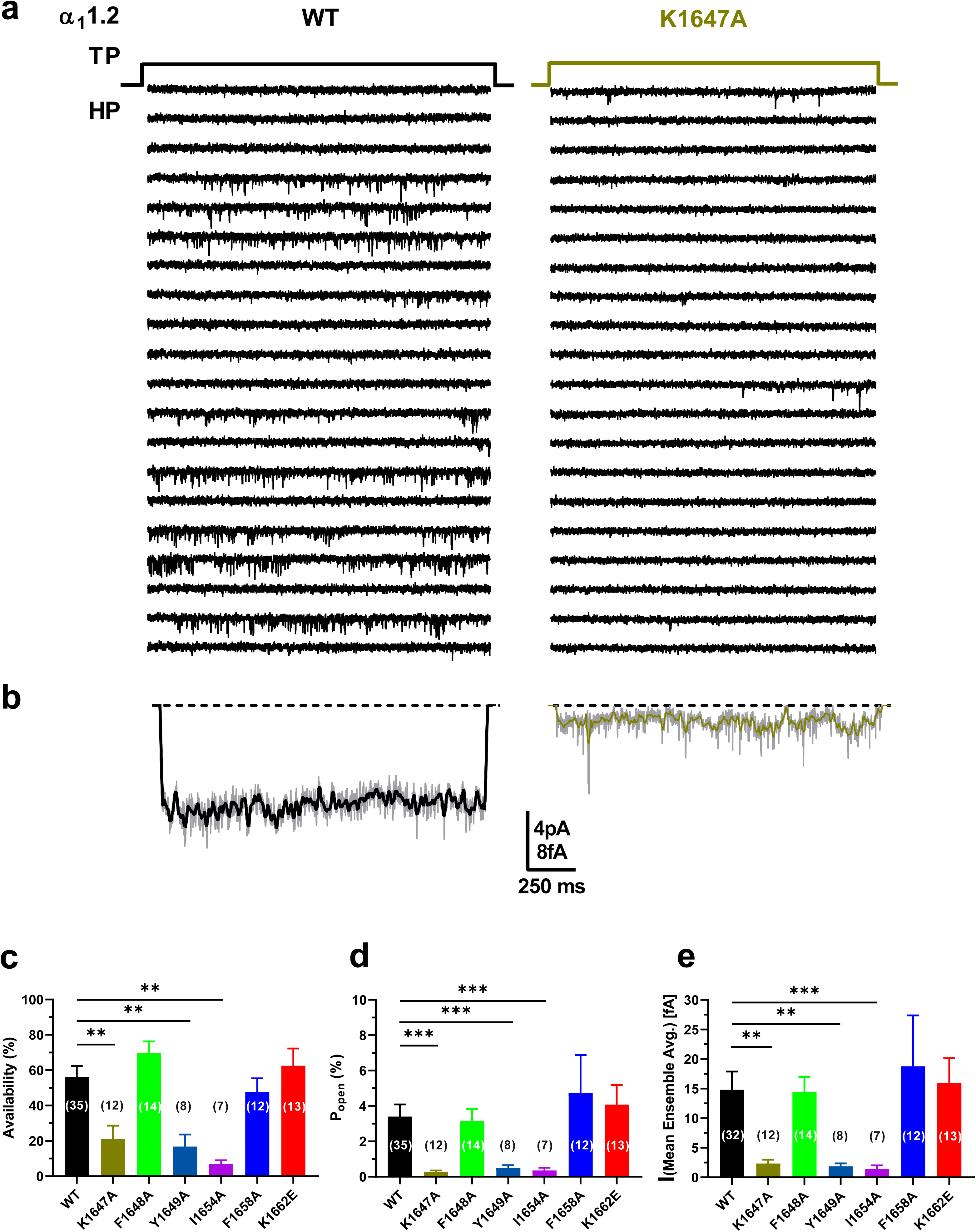
Ca_V_1.2 mutations that affect α-Actinin-1 drastically decrease channel open probability. HEK293 cells were transfected with α_1_1.2, α_2_δ-1, and β_2A_ and cultured for 22-24h before cell attached patch recording in 110 mM Ba^2+^. a, Single Ca_V_1.2 channel recordings of Ca_V_1.2 WT and K1647A. Holding potential (HP) was −80 mV and test potential (TP) 0 mV. Shown are 20 consecutive sweeps from representative experiments. b, Mean assembly averages for all experiments with Ca_V_1.2 WT and K1647A, which are based on a total of 2744 and 868 sweeps, respectively (see Table 5). c-e, Means ± SEM for availability (i.e., likelihood that a sweep had at least one event; c), Po (d), and the mean of the current of a single channel at any point in time calculated from the ensemble averages of each experiment (e) (**p<0.01, ***p<0.001 compared to WT; one-way ANOVA with Bonferroni post-hoc test (c) or Welch ANOVA with Tamhane T2 test (d,e); n = 7 to 35 (given in parenthesis in each column); see Suppl. Fig. 5, 6 and Table 5 for more details).

**Table 5:**
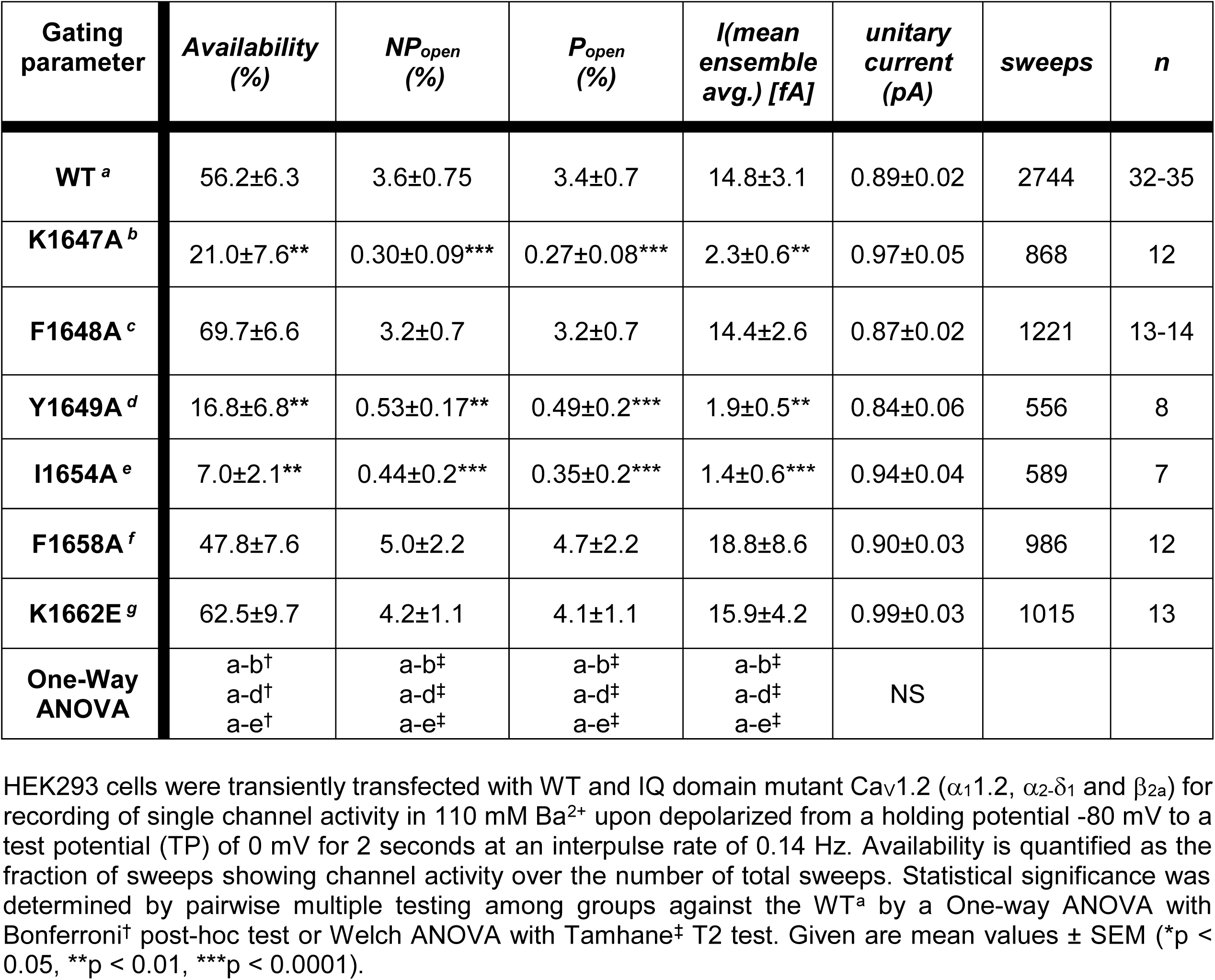
Biophysical properties of Ca_V_1.2 microscopic single channel currents for WT versus IQ motif mutants

The single channel activity is the product of the number (N) of channels in a patch, the open probability (Po), and unitary current amplitudes (i) of each individual channel (i.e., N·Po·i). Because i was unaffected (Suppl. Fig. 6e) the mutations must affect NPo. Given that surface expression of all three mutations decreased by only 35-40% versus WT α_1_1.2 under exactly the same conditions (Tseng et al., 2017), the ∼90% reduction of NPo for all three mutants to ∼10% of WT Ca_V_1.2 suggests that Po of the remaining ∼60% channels is only ∼ 1/6 of the Po of WT Ca_V_1.2. These observations suggest a remarkable ∼6-fold decrease in Po upon loss of α-actinin binding. To corroborate this notion, we determined N for each recording and derived Po for individual channels. Accordingly, Po is reduced by ∼90% for individual channels carrying a K1647A, Y1649A, or I1654A mutation (Fig. 3d; Suppl. Fig. 6c; Table 5). At the same time, the F1658A and K1662E mutations, which diminish apoCaM binding, did not alter NPo.

To further scrutinize the role of α-actinin in Po of Ca_V_1.2, Ca_V_1.2 was co-expressed with WT α-actinin-1 or its binding-deficient E847K/E851K mutant. Overexpression of WT but not E847K/E851K mutant α-actinin-1 increased Po of Ca_V_1.2 WT >2-fold from 3.4% seen with Ca_V_1.2 alone to 7.9% (Fig. 4a,d; Suppl. Fig. 7c; Table 6). Availability, NPo, and assemble averages were also increased (Fig. 4b,c,e; Suppl. Fig. 7a,b,d; Table 6). Accordingly, in HEK293 cells functional occupancy of Ca_V_1.2 by endogenous α-actinin appears to be far from complete.

**Fig. 4.**
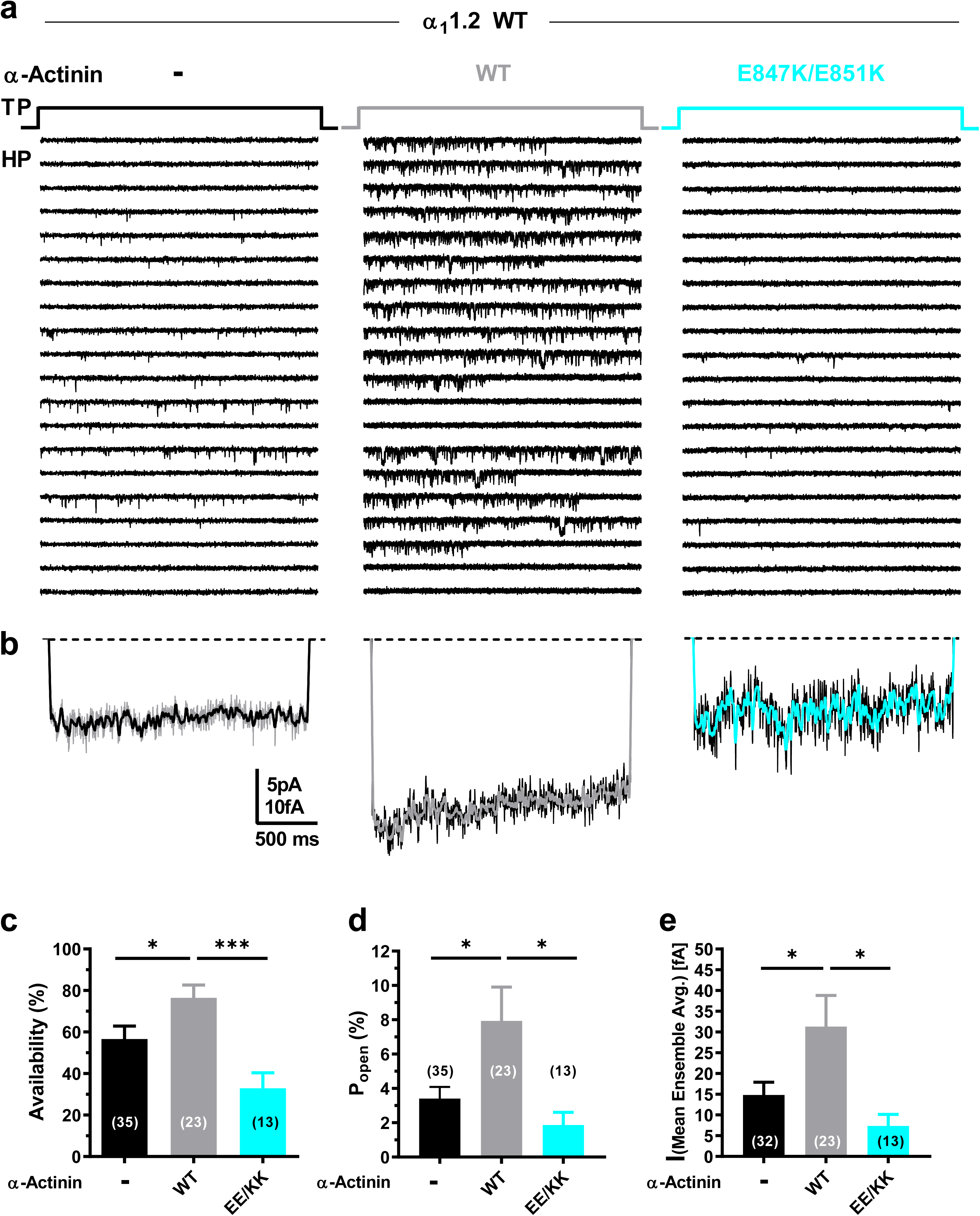
Ectopic α-actinin-1 expression increases Ca_V_1.2 open probability. HEK293 cells were transfected with α_1_1.2, α_2_δ-1, and β_2A_ plus, if indicated, α-actinin-1 before cell attached patch recording. a, Representative single Ca_V_1.2 channel recordings of WT Ca_V_1.2 alone or with WT or E847K/E851K mutant α-actinin-1. b, Mean assembly averages for all experiments for each combination. c-e, Means ± SEM for availability (c), Po (d), and the mean of the current of a single channel calculated from the ensemble averages of each experiment (e) (*p<0.05, **p<0.01, ***p<0.001; unpaired t-test; n = 13 to 35; see Suppl. Fig. 7 and Table 6 for more details).

**Table 6:**
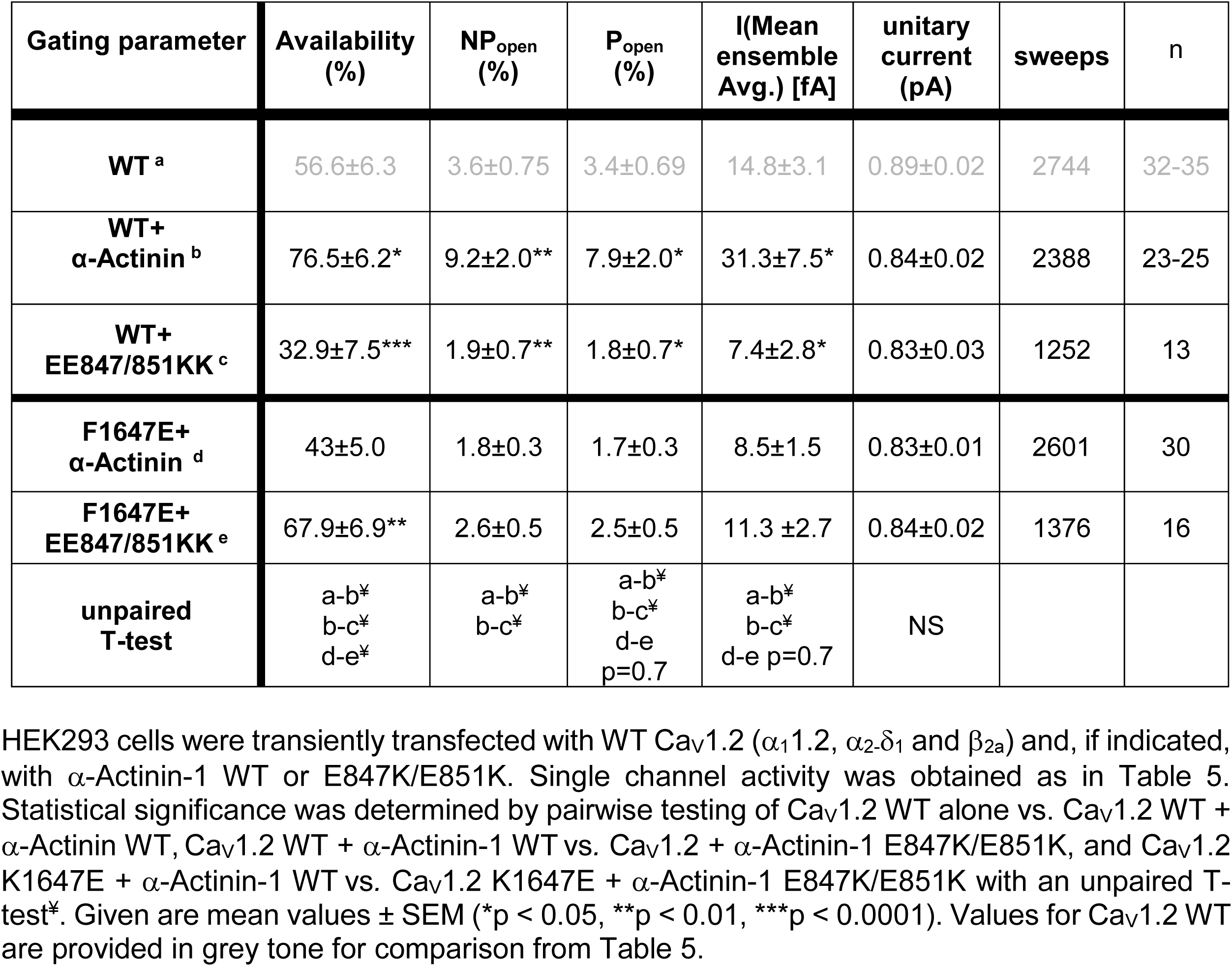
Effect of α-actinin-1 ectopic expression on biophysical properties of Ca_V_1.2 single channel currents

To further test whether α-actinin needs to bind to the Ca_V_1.2 IQ motif to augment Po, we used a charge inversion experiment by mutating K1647 to the negatively charged glutamyl rather than neutral alanyl residue and then attempted rescue of the expected reduction in Po by pairing expression of K1647E Ca_V_1.2 with charge inverted E847K/E851K α-actinin-1. Coexpression of K1647E mutant Ca_V_1.2 with WT α-actinin-1 yielded a low availability, NPo, Po, and ensemble average (Fig. 5; Suppl. Fig. 8; Table 6), which were well below the values observed upon expression of WT Ca_V_1.2 alone or co-expression of WT Ca_V_1.2 with E847K/E851K mutant α-actinin-1 (Fig. 4; Table 6). The reduction in availability was statistically significantly, though not fully, rescued when K1647E mutant Ca_V_1.2 was paired with E847K/E851K mutant rather than WT α-actinin-1 (Fig. 5a; Suppl. Fig. 8a; Table 6). Po also appears to be partially rescued although the p value was 0.07 versus pairing of WT α-actinin-1 with K1647E mutant Ca_V_1.2.

**Fig. 5.**
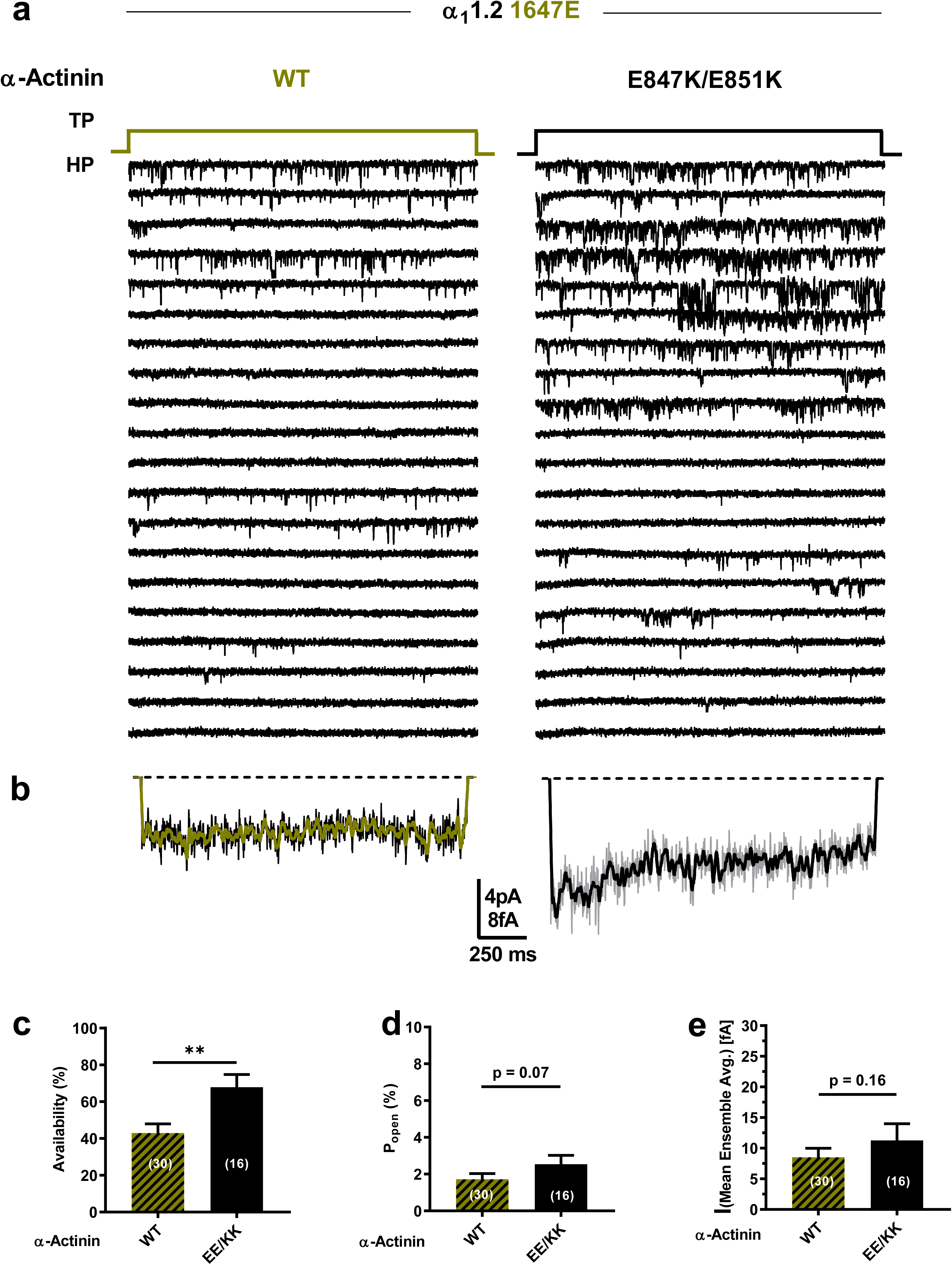
α-Actinin-1 E847K/E851K rescues open probability for Ca_V_1.2 K1647E. HEK293 cells were transfected with α_1_1.2 with the K1647E mutation, α_2_δ-1, and β_2A_ plus α-actinin-1 WT or E847K/E851K before cell attached patch recording. a, Representative single Ca_V_1.2 channel recordings of Ca_V_1.2 K1647E with WT or E847K/E851K α-actinin-1. b, Mean assembly averages for all experiments for both combinations. c-e, Means ± SEM for availability (c), Po (d), and mean of the current of a single channel calculated from the ensemble averages of each experiment (e) (**p<0.01; unpaired t-test; n = 16 to 35=0; see Suppl. Fig. 8 and Table 6 for more details).

This rescue of loss of availability and likely of Po for K1647E mutant Ca_V_1.2 by E847K/E851K mutant α-actinin-1 must be due to either enhanced opening of individual channels, a change in channel surface expression, or both. This rescue is difficult to explain by a mechanism other than that α-actinin-1 binding augments channel activity, including apoCaM binding. Consistently, impairing apoCaM binding to the Ca_V_1.2 IQ motif by mutating F1658 to Ala or K1662 to Glu had no effect on availability, NPo, Po, and did not affect ensemble averages (Fig. 3; Suppl. Fig. 5, 6).

One possibility for a reduction in Po is that K1647A, Y1649A, and I1654A shift the voltage dependence of Ca_V_1.2 to more positive potentials such that WT channels open upon depolarization to 0 mV more readily than mutant channels. However, neither the reversal potential nor the voltage dependence of activation was significantly affected by the K1647A and Y1649A mutations and only minimally by the I1654A mutation (Suppl. Fig. 9B,C; Table 7).

**Table 7.** Biophysical properties of Ca_V_1.2 macroscopic whole cell patch currents for WT versus IQ motif mutants

| Analysis | I-V | | | G-V | | | Charge movement | $I_{\text{Tail}}/Q_{\text{on}}$ |
| --- | --- | --- | --- | --- | --- | --- | --- | --- |
| Parameter | $V_{1/2\text{act.}}$<br>(mV) | $k_{\text{act.}}$ | $E_{\text{rev.}}$ (mV) | $V_{1/2\text{act.}}$<br>(mV) | $k_{\text{low}}$ | $k_{\text{high}}$ | $Q_{\text{on}}$<br>(fC/pF) | slope |
| WT<sup>a</sup> | -18.5±0.5 | -4.2±0.4<br>(10) | 51.3±1.3 | -6.5.0±0.4 | 7.0±0.3<br>(16) | -8.5±27.4 | 3.8±0.9<br>(18) | 3.9±1.5<br>(16) |
| K1647A<sup>b</sup> | -19.9±0.9 | -5.9±0.6<br>(11) | 55.2±2.0 | -8.5.0±0.9 | 8.5±0.8<br>(15) | 9.0±10.5 | 1.3±0.3*<br>(15) | 0.26±1.5<br>(15) |
| Y1649A<sup>c</sup> | -19.9±0.6 | -4.0±0.5<br>(10) | 49.2±1.5 | -6.3.0±0.7 | 7.5±0.8<br>(16) | 28.7±6.9 | 0.98±0.3**<br>(13) | 0.19±1.0<br>(12) |
| I1654A<sup>d</sup> | -12.6±1.0**** | -5.4±0.7<br>(5) | 48.2±2.6 | -2.5±0.9** | 7.5±0.9<br>(13) | 38.5±11.5 | 0.87±0.3**<br>(13) | 1.3±0.6<br>(12) |
| F1658A<sup>e</sup> | -26.5±1.1** | -4.5±0.9<br>(6) | 58.6±4.1 | -10.6±1.4** | 5.3±1.6<br>(11) | 20.2±3.9 | 2.8±0.6<br>(13) | 3.1±1.4<br>(12) |
| K1662E<sup>f</sup> | -20.2±0.4 | -4.2±0.3<br>(12) | 50.1±1.2 | -6.1.0±0.7 | 7.9±0.8<br>(21) | 49.2±19.4 | 3.5±1.1<br>(17) | 2.2±1.0<br>(14) |
| ANOVA/<br>T-Test | a-d<sup>†</sup><br>a-e<sup>†</sup> | NS | NS | a-d<sup>†</sup><br>a-e<sup>†</sup> | NS | NS | a-b<sup>†</sup><br>a-c<sup>†</sup><br>a-d<sup>†</sup> |  |

To test whether this charge-inversion rescue for the K1647E mutant promoted surface expression of Ca_V_1.2 we performed surface biotinylation experiments (Fig. 6). As expected, co-expression of WT α-actinin-1 with WT Ca_V_1.2 increased α_1_1.2 biotinylation by ∼60% versus WT α_1_1.2 alone. Thus, WT Ca_V_1.2 expression at the surface is enhanced by α-actinin-1 (Fig. 6a). That the increase in surface expression of WT Ca_V_1.2 by WT α-actinin-1 overexpression was smaller than the increase in Po is analogous to the smaller effects of the α_1_1.2 K1647A, Y1649A, and I1654A mutations on surface expression compared to charge density (Tseng et al., 2017) and the larger decrease in Po and NPo we report here (Fig. 3). Importantly, this increase in Ca_V_1.2 surface expression by α-actinin-1 was lost when WT α_1_1.2 was co-expressed with E847K/E851K mutant α-actinin-1 or K1647E mutant α_1_1.2 with WT α-actinin-1 (Fig. 6b). However, the charge inversion we performed by combining these mutants failed to increase surface expression of Ca_V_1.2 to a degree that would be detectable (Fig. 6b). This result is in contrast to our findings that K1647E α_1_1.2 availability and likely Po was partially rescued by the E847K/E851K α-actinin-1 charge reversal (Fig. 5). It is conceivable that a small rescue effect did occur but was indiscernible for statistical reasons as 95% confidence intervals (CIs) are larger than a potentially partial rescue of, e.g., 30% of the impairment of ∼50% by the K1647E mismatch with WT α-actinin-1 (Suppl. Table 1; a 30% rescue would translate into only an ∼15% increase in surface expression).

**Fig. 6.**
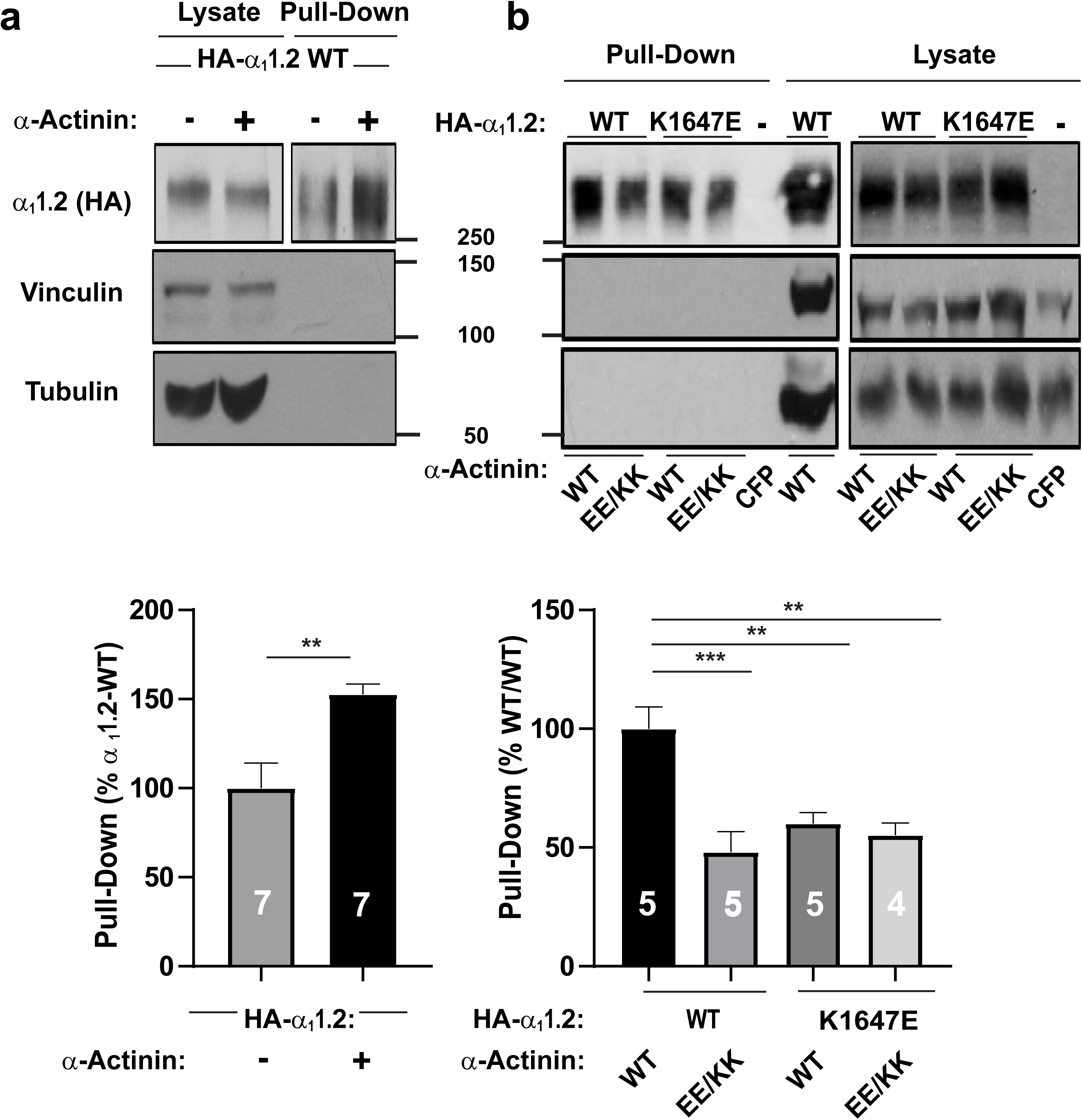
Modulation of Ca_V_1.2 surface expression via its interaction with α-actinin-1. a, HEK293 cells were transfected with WT α_1_1.2, α_2_δ-1, and β_2A_ ± WT α-actinin-1 and cultured for 22-24h before surface biotinylation. Shown are representative immunoblots of NeutrAvidin pull-down samples (from lysate containing 600 μg protein) and total lysate samples containing 20 μg protein. α_1_1.2 was detected with an antibody against its HA tag (*top*), which is present in all constructs used throughout this work. Pull-down and lysate samples are from the same blot but different exposures because signals from lysate samples were much stronger than from pull-down samples. Immunoblotting for vinculin (*middle*) and tubulin (*bottom*) indicated that comparable amounts of these intracellular control proteins were present in lysate samples. Their absence in pull-down samples as seen on the same blots showed that these prominent intracellular proteins did not undergo biotinylation as control for membrane integrity during surface biotinylation. Bar graph shows means ± SEM of the pull-down immunosignals in mutants relative to surface labelling of control α_1_1.2 samples lacking α-actinin-1 co-expression (mean set to 100%; see Methods; *p<0.05; two-tailed t-test; n = 7). b, HEK293 cells were transfected with WT or K1647E mutant α_1_1.2, α_2_δ-1, and β_2A,_ and WT or E847K/E851K mutant α-actinin-1, or CFP alone as negative control and cultured for 22-24h before surface biotinylation. Shown are representative immunoblots of pull-down and lysate samples as in (a). Bar graph shows means ± SEM of the pull-down immunosignals in mutants relative to surface labelling in the WT/WT control (mean set to 100%; see Methods). **p<0.01, ***p < 0.001; one-way ANOVA with Tukey post hoc test; n = 4-5).

### α-Actinin binding to the Ca_V_1.2 IQ motif augments gating charge movement as well as its coupling to channel opening

To test the molecular mechanism underlying the α-actinin-induced increase in Po in more detail we analyzed gating charge movement at the Ba^2+^ current reversal potential (Fig. 7) by whole-cell patch clamp recording. This approach measures all of the Ca^2+^ channels on the surface, thereby ruling out any potential limitation caused by the channel selection inherent in the cell-attached patch recordings in Figs. 3-5. The K1647A, Y1649A, and I1654A mutations reduced gating currents (Q_on_) by 66-77% (Fig. 7a,b,d; Suppl. Fig. 9a; Table 7) suggesting that α-actinin fosters gating charge movement. This reduction is stronger than the reduction in surface expression (Tseng et al., 2017) but not quite as strong as the 90% reduction in NPo for these mutants (Fig. 5). Thus, we investigated whether α-actinin also augments coupling of gating charge movement to channel opening. We analyzed the relationship of tail currents (I_tail_) to Q_on_ and determined the slopes of the regression curves as an established parameter for quantification of the coupling of Q_on_ to I_tail_ (Tuluc et al., 2009, Yang et al., 2010). Accordingly, the slopes for K1647A, Y1649A, and I1654A mutant Ca_V_1.2 were strongly reduced compared to WT (Fig. 7c; Table 7). Because large gating events were rare for these mutants, we combined all data from those three mutants for further analysis to better cover that range of the curve. This analysis confirmed that the slope of the resulting curve was substantially smaller than for WT Ca_V_1.2 (Fig. 7e). Collectively, these data indicate that binding of α-actinin to the Ca_V_1.2 IQ motif augments not only surface expression but also gating charge movement and its coupling to channel opening. Neither gating charge movements nor their coupling to channel opening were significantly affected by the F1658A or K1662E mutations (Fig. 7; Suppl. Fig. 9a, Table 7), once more arguing against a role in apoCaM binding to the IQ motif in defining NPo under basal conditions.

**Fig. 7.**
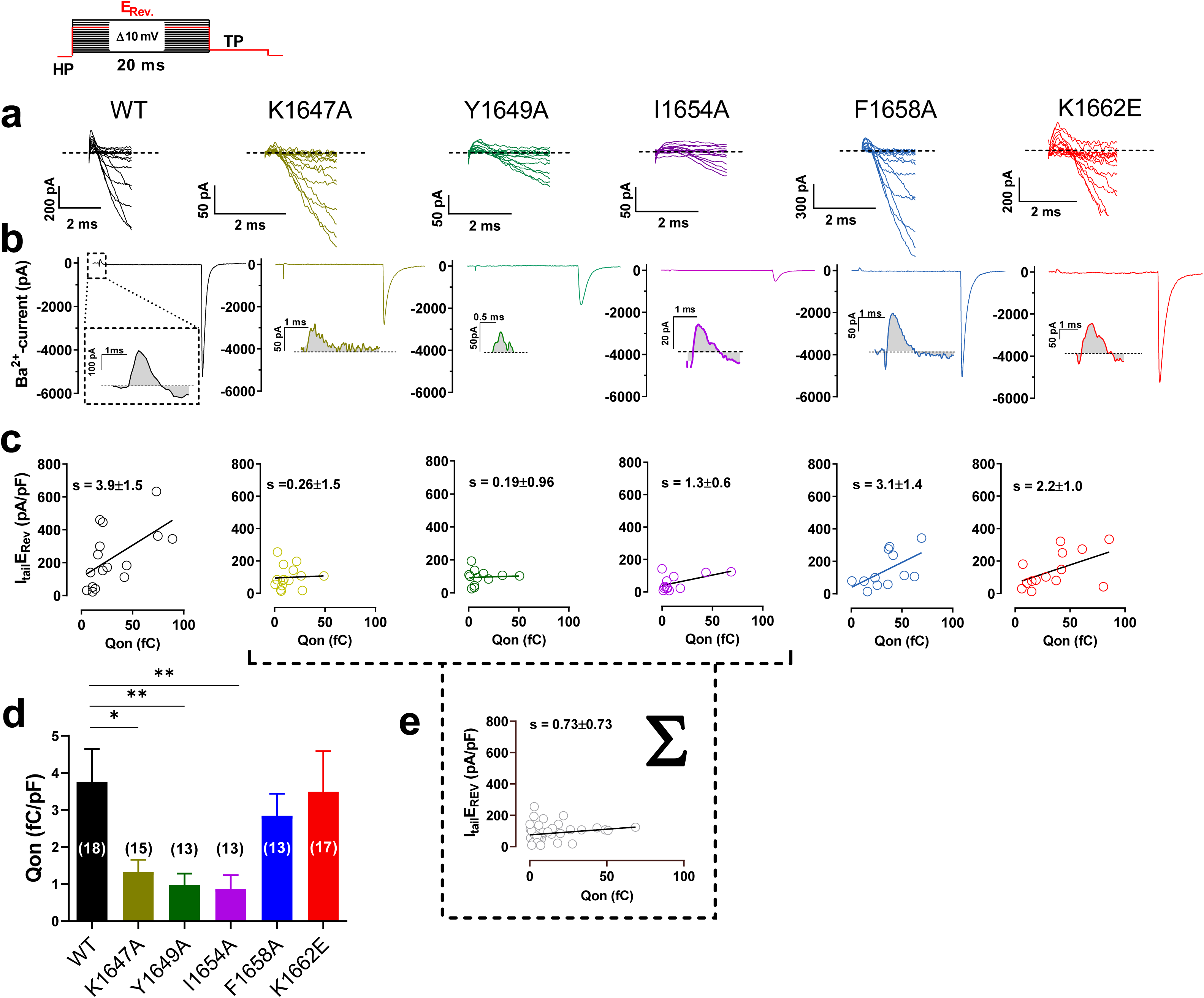
Ca_V_1.2 mutations that affect α-actinin-1 impair gating charge movement and its coupling to channel opening. HEK293 cells were transfected with α_1_1.2, α_2_δ-1, and β_2A_ before whole cell patch recording in 20 mM Ba^2+^. a, Representative current traces of the first 2 ms obtained from recordings upon depolarizations from a holding potential of −80 mV to the indicated potentials (the voltage protocol is schematized in the upper left corner). b, Representative current traces upon step depolarizations to the reversal potential (E_rev_) for 20 ms to obtain movement of the ON-gating charges (Q_on_), and subsequent to −50 mV for 10 ms to obtain tail currents (I_tail_). Insets: magnifications of exemplary Q_on_ for Ca_V_1.2 WT, K1647A, Y1649A, I1654A, F1658A and K1662E. c, Plots of I_tail_ (in this panel corrected for variations in cell capacitance) versus total detectable charge transfer for Q_on_. Slopes of regression curves are strongly reduced for Ca_V_1.2 K1647A, Y1649A and I1654A versus WT (see Table 7 for more details). d, Means ± SEM of Q_on_ (in this panel corrected for variations in cell capacitance; *p<0.05; **p<0.01; one-way ANOVA with Bonferroni post-hoc test; n = 13-18; see Suppl. Fig. 9 and Table 7 for more details). e, The reduction in slope of the regression curve of combined population data for K1647A, Y1649A and I1654A versus WT indicates reduced coupling of I_tail_ with Q_on_ when α-actinin binding to the IQ motif is diminished.

## DISCUSSION

The NMR structures of α-actinin-1 and apoCaM bound to the Ca_V_1.2 IQ motif presented here revealed distinct intermolecular interactions that were verified by mutagenesis experiments involving charge inversion for both interactions. The EF hands in α-actinin-1 are not capable of binding to Ca^2+^ under physiological conditions because their EF-hand loops lack the proper residues for ligating Ca^2+^ with high affinity (Backman, 2015). Therefore, the α-actinin-1 EF-hand domain bound to Ca_V_1.2 is in the Ca^2+^-free state under basal conditions ([Ca^2+^]_i_ of ∼100 nM) and adopts a semi-open conformation of Ca^2+^-free EF-hands like that observed for apoCaM bound to various target proteins (Chagot and Chazin, 2011, Feldkamp et al., 2011, Houdusse et al., 2006). However, the semi-open EF-hand conformation of α-actinin-1 binds to the IQ peptide with a polarity opposite to that of apoCaM (see black arrows in Suppl. Fig. 10a,b). The Ca_V_1.2 residue K1647 at the N-terminal end of the IQ helix forms intermolecular salt bridges with α-actinin-1 residues E847 and E851. By stark contrast, the IQ helix is rotated 180 degrees upon binding to apoCaM (see black arrows in Suppl. Fig. 10). This opposite orientation places Ca_V_1.2 residue K1662 at the C-terminal end of the IQ helix in close proximity to CaM residues E85 and E88, which are homologous to α-actinin-1 residues E847 and E851. Similarly, the IQ helices in voltage-gated Na^+^ channels bind to apoCaM and Ca^2+^-saturated CaM with opposite polarity (Hovey et al., 2017). Thus, the orientation of the IQ helix bound to the EF-hand motif may be an important structural determinant of binding specificity and may explain why α-actinin-1 and apoCaM have different functional effects.

Voltage-gated Na^+^ channels exhibit Ca^2+^-dependent inactivation (CDI) mediated by Ca^2+^-CaM (Ben-Johny et al., 2014, Gabelli et al., 2016), similar to Ca_V_1.2 (Peterson et al., 1999, Zuhlke et al., 1999). However, the structure of apoCaM bound to the Ca_V_1.2 IQ motif (Suppl. Fig. 10b) is quite different from previous structures of apoCaM bound to Na_V_1.2 (Suppl. Fig. 10c) or Na_V_1.5 (Chagot and Chazin, 2011, Gabelli et al., 2014). The Na_V_1.2 IQ motif sequence is only 17% identical to that of Ca_V_1.2 (Suppl. Fig. 10d). Na_V_1.2 residues A1909, A1915, and Y1919, which are not conserved in Ca_V_1.2, each make critical contacts with apoCaM. (Suppl. Fig. 10c). The side chain methyl groups of A1909 and A1915 are each 2.5 Å away from the side chain methyl atoms of CaM residues M109 and L85, respectively. The aromatic side chain of Y1919 is in intimate contact with the aromatic side chain of F141 from CaM. Another important difference is that the Ca_V_1.2 IQ helix binds to apoCaM with opposite directionality compared to the Na_V_1.2 IQ helix (see black arrows in Suppl. Fig. 10b,c). Hence, the salt bridge that connects Ca_V_1.2 (K1662) to apoCaM (E88) is not conserved in Na_V_1.2 or Na_V_1.5 but the large number of non-conserved intermolecular hydrophobic contacts to Na_V_1.2 (Suppl. Fig. 10c) caused by the opposite binding orientation of the IQ helix should outweigh any stabilization from the salt bridge in Ca_V_1.2 and may explain why apoCaM binds to Na_V_1.2 and Na_V_1.5 with nanomolar affinity (Hovey et al., 2017) compared to the much weaker micromolar affinity of apoCaM binding to Ca_V_1.2 (Table 4).

The relatively high dissociation constant for apoCaM binding to the α_1_1.2 IQ motif (K_d_ = 10 μM) may be outside the physiological concentration range for apoCaM in neurons. Under basal conditions, the cytosolic apoCaM is kept below 1μM (Wu and Bers, 2007) by proteins that have a high affinity for apoCaM. For instance, in neurons neurogranin (RC3) (Huang et al., 2004, Huang et al., 2000, Ran et al., 2003, Zhong et al., 2011) and GAP-43 (P-57) (Cimler et al., 1985) serve as sinks and reservoirs for apoCaM. In neurons, total concentration of CaM is ∼10 μM (Egrie et al., 1977, Zhabotinsky et al., 2006), of neurogranin is 20 μM (Huang et al., 2004, Zhabotinsky et al., 2006), and of GAP-43 is 18 μM (Cimler et al., 1985, Zhabotinsky et al., 2006), and their Kd values for apoCaM are in the range of 1-5 μM (Alexander et al., 1987, Huang et al., 2000, Zhabotinsky et al., 2006), resulting in 0.9 μM free CaM under basal conditions (Zhabotinsky et al., 2006). Therefore, on the basis of the relatively low binding affinity of apoCaM (K_d_ = 10 μM), the fractional binding of apoCaM to α_1_1.2 can be estimated to be less than 10% 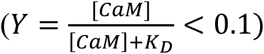 under basal physiological conditions in neurons. Recent work suggests that ectopically expressed WT CaM as well as Ca^2+^ binding-deficient CaM_1234_ is in large excess over endogenous CaM (Iacobucci and Popescu, 2017). As a result, CaM_1234_ may occupy a much larger fraction of α_1_1.2 than the endogenous WT apoCaM. In this scenario expression of CaM_1234_ could impair CDI as seen earlier (Peterson et al., 1999) by mechanisms other than displacement of endogenous apoCaM as invoked earlier (Peterson et al., 1999).

Alterations of I1609 in α_1_1.3 (equivalent to I1654 in α_1_1.2) impair binding of apoCaM to the closely related Ca_V_1.3 channel and Po, which was interpreted to mean that apoCaM binding to this IQ motif augments Po (Adams et al., 2014). Here we show that the homologous I1654 in the highly conserved IQ motif of Ca_V_1.2 is not only critical for apoCaM but also α-actinin binding. Accordingly, mutating I1609 could have decreased NPo by impairing α-actinin rather than apoCaM binding. Indeed, cell attached channel recordings showed an ∼90% reduction in Po for all three α-actinin binding - deficient α_1_1.2 IQ mutants. This effect included K1647, which contacts α-actinin (Fig. 1c) but not apoCaM. Consistently, the K1647A mutation reduced binding of α-actinin (Table 2) but not apoCaM (Table 4). The same is true for the Y1649A mutation. Although Y1649 does not form a direct interaction with α-actinin EF3-EF4, the Y1649A mutation does abrogate α-actinin binding in the yeast two hybrid system (Tseng et al., 2017). We suggest that Y1649 is necessary to stabilize the orientation of L1653 and neighboring I1654, which is required for binding to α-actinin (Suppl. Fig. 11a). Thus, our new data indicate that Po is largely determined by α-actinin and not apoCaM binding. In fact, the F1658A and K1662E mutations, which decreased apoCaM binding (Table 4), did affected neither Po (Fig. 3e) nor Q_on_ (Fig. 7a,b,d) and minimally if at all α-actinin binding (Table 2). In support of these results, others found no effect of CaM on Ca_V_1.2 plasma membrane targeting in HEK293 cells (Bourdin et al., 2010). This conclusion is further supported by the charge inversion experiments of K1647E mutant α_1_1.2 and E847K/E851K mutant α-actinin-1 in which impaired functional channel availability and likely impaired Po is partially rescued (Fig. 5). These findings support the functional relevance of the interaction of the positively charge K1647 in the IQ motif with the negatively charged E847 and E851 of α-actinin-1. That the rescue of the reduction in Po of K1647E mutant α_1_1.2 by E847K/E851K mutant α-actinin-1 is far from complete can in part be explained by analysis of the E851K rotamers (Suppl. Fig. 11b-d). Most of the E851K rotamers are predicted to clash with side chain and backbone atoms within the α-actinin region at the channel - α-actinin interface based on our structure (Fig. 1c), which will most likely affect the positioning of the EF hand region of α-actinin relative to α_1_1.2 and thereby the α-actinin - α_1_1.2 interaction. As a result, binding of E847K/E851K mutant α-actinin-1 to α_1_1.2 will be decreased as compared to WT α-actinin. In fact, binding affinity is also not fully rescued when the K1647E IQ peptide is titrated with E847K/E851K mutant α-actinin-1 EF3/4 segment (Table 2). A change in the exact structure of the region surrounding the IQ motif in full length α_1_1.2 could be the reason for further impairment of α-actinin binding furnishing a potential explanation for our finding that functional availability and Po is less effectively rescued by the full charge inversion than the in vitro binding affinity.

Further mechanistic insight is provided by the strong reduction in gating charge movement upon loss of α-actinin binding (Fig. 7). The most striking and very clear finding for Q_on_ is the observation that gating charges for the three α-actinin binding-deficient α_1_1.2 mutants are so small that they are often barely if at all detectable and amplitudes are difficult to determine. Accordingly, α-actinin binding to the IQ motif promotes the outward movement of the S4 segments, which gives rise to the gating charges, presumably by either lowering energy barriers for this motion or by stabilizing the ‘outward’ conformation of S4 segments. In addition, α-actinin binding to the IQ motif augments also the coupling between this gating movement and channel opening. This finding is reminiscent of the effect of Ca_V_1.2 phosphorylation by PKA on S1700, which upregulates this coupling (Fuller et al., 2010, Fuller et al., 2014). Because of the close proximity of S1700 to the IQ motif it is tempting to speculate that phosphorylation of S1700 and α-actinin binding to the IQ motif intersect to facilitate channel gating, possibly through similar effects on overall conformations of Ca_V_1.2.

All three α-actinin binding - deficient IQ mutants of Ca_V_1.2 reduced NPo by ∼ 90% when channel density was reduced by only 35-40% as seen by surface labeling (Tseng et al., 2017). To further ascertain that a large portion of the reduction in NPo is due to a reduction in Po and not just N, i.e., not only due to a reduction in surface expression, we determined the 95% confidence intervals (CI) for surface biotinylation, surface labeling with antibodies with subsequent analysis by fluorescence activated cell sorting (Bourdin et al., 2010, Yang et al., 2010), Po, and Q_on_ (Suppl. Tables 1-3). The CIs for surface biotinylation and surface antibody labeling are remarkably similar (e.g., 52-69% and 51-71%, respectively, for the most relevant mutant K1647A, with WT being 100%) and overlap not at all with Q_on_ and Po (26-44% and 6-10%, respectively, for K1647A, with WT again being 100%). Thus, the CIs for the 95% CIs for reductions are ∼29-49% for surface stainings, 56-74% for Q_on_, and 90-94% for Po. Accordingly, it appears extremely unlikely that the reductions in surface expression can fully account for the reductions in Q_on_ and Po for α-actinin – binding-deficient Cav1.2; rather loss of surface localization is only responsible for a fraction of the reductions in Q_on_ and Po. Similarly, the reduction in Q_on_ can most likely only partially account for the reduction in Po, supporting the notion that coupling between the movement of the voltage sensor and the channel gate is impaired in addition to the movement of this sensor.

If loss of α-actinin binding to the IQ motif would not affect Po of individual channels, then NPo of all α-actinin binding - deficient IQ mutants should be ∼60% of wild type Ca_V_1.2. A 90% decrease in NPo means that 60% of the Ca_V_1.2 channels that remain on the cell surface upon impairment of α-actinin binding carry only ∼10% of the current seen for the wild type Ca_V_1.2 population. These effects translate into a 6-fold reduction in Po of individual Ca_V_1.2 channels upon loss of α-actinin binding.

Together with earlier work that showed that α-actinin increases surface localization and postsynaptic accumulation of Ca_V_1.2 (Hall et al., 2013, Tseng et al., 2017), this new work now demonstrates that α-actinin serves a dual role in regulating Ca_V_1.2 function. It not only promotes Ca_V_1.2 surface expression but, remarkably, also exerts a strong positive effect on Po. This mechanism allows Ca_V_1.2 to be minimally active during secretory trafficking or when it is outside its target regions but to become functionally fully engaged when anchored at proper locations such as postsynaptic sites. Thus, coupling of α-actinin binding to both localization and activity of Ca_V_1.2 are perfectly fine tuned. This mechanism is so far unique. Whether analogous mechanisms apply to other channels and especially other Ca^2+^ channels, which may require potent mechanisms to prevent inappropriate and potentially harmful Ca^2+^ flux from secretory compartments or at the wrong locations on the cell surface, is now an intriguing premise that will inspire future work.

## MATERIALS AND METHODS

### NMR spectroscopy

Xenopus calmodulin was expressed in *E. coli* strain BL21(DE3) grown in LB medium (unlabeled proteins) or M9 media supplemented with ^15^NH_4_Cl or ^15^NH_4_Cl/^13^C-glucose for single- or double-labeled proteins, respectively. Recombinant CaM was prepared as described previously(Zhang et al., 2012).

Human α-actinin-1_EF12 (residues 750-812) and α-actinin-1_EF34 (residues 822-893) were each subcloned into pET3b vector and expressed in *E. coli* strain BL21(DE3) grown in LB medium (unlabeled proteins) or M9 media supplemented with ^15^NH_4_Cl or ^15^NH_4_Cl/^13^C-glucose for single- or double-labeled proteins, respectively. α-actinin-1_EF12 and α-actinin-1_EF34 were each purified and prepared as described (Turner et al., 2016).

Unlabeled Ca_V_1.2 IQ peptide (residues 1644-1664) was purchased from ChinaPeptides. The peptide was dissolved in d6-DMSO to give a peptide concentration of 7.8 mM. An aliquot of peptide (1.5 equivalents) was added to a dilute solution of α-actinin-1_EF34 or apoCaM (50 μM protein dissolved in 20 mM 2-Amino-2-hydroxymethyl-propane-1,3-diol-d11 (Tris-*d*_11_) with 50 mM NaCl, 5mM dithiothreitol-d10 (DTT-*d10)* and 95% H_2_O/5% D_2_O) and incubated at 15^0^C for 1 hour to ensure complete binding of the peptide. The complex was then concentrated to a final concentration of 500 μM in a final volume of 500 μL for NMR experiments.

### Production and use of isotope-labeled peptides using pET-31

We expressed ^15^N- or ^15^N/^13^C-labeled IQ peptide in *E. coli* as a fusion protein with the ketosteroid isomerase (KSI) using the pET31 plasmid (Kuliopulos et al., 1994) (Novagen/EMD Biosciences). KSI fusion proteins are concentrated in inclusion bodies in *E. coli*, protecting the fused peptide from proteolysis. The fusion protein was purified by affinity chromatography under denaturing conditions via a 6x-His tag at the C-terminus. To release the peptide of interest, the purified fusion is cleaved using cyanogen bromide (CNBr). The CNBr cleaves at methionine residues that were engineered between the KSI and peptide, and between the peptide and 6x-His tag. This cleavage mixture is then rotary evaporated to dryness and resuspended in an appropriate ratio of acetonitrile/water, which solublizes primarily the peptide and not KSI. Finally, the peptide is purified by reverse-phase HPLC.

Two complementary oligonucleotides that code for Ca_v_1.2 IQ peptide (5’p-TTT TAT GCG ACC TTT CTG ATT CAG GAA TAT TTT CGC AAA TTT AAA AAA CGC AAA <u>ATG</u>; 5’p-TTT GCG TTT TTT AAA TTT GCG AAA ATA TTC CTG AAT CAG AAA GGT CGC ATA AAA <u>CAT</u>) were annealed in 40 mM Tris-HCl, pH 8.0, 10mM MgCl_2_, 50 mM NaCl by heating to 95° C and then cooling slowly (>2h) to room temperature (RT). The annealed, double-stranded insert was then ligated into AlwNI-digested and dephosphorylated pET31 using T4 DNA Ligase. The 5’-phosphorylation augments ligation and the extra Met codons (underlined) provide compatible sticky ends for the AlwNI-cut plasmid as well as the sites of CNBr cleavage. 2 ul of the ligation mixture were transformed into DH5alpha cells and plasmids from individual ampicillin-resistant colonies sequenced. Successful insertions were obtained, including some multiple insertion. We used one of the single-insertion plasmids for expression in M9 minimal media. *E. coli* was lysed by sonication. The insoluble material was collected by ultracentrifugation and resuspended in 6M guanidine, insoluble material removed by ultracentrifugation, and the supernatant loaded onto a Ni-NTA column equilibrated with 6M guanidine buffer. After elution with 300 mM imidazole, the purified fractions were pooled and then dialyzed against ultrapure water, causing the fusion protein to precipitate. Precipitate was collected by centrifugation and resuspended in 80% formic acid, transferred to a round-bottom flask and injected with N_2_ at ∼3 psi. For a 2L scale expression, we used 60 ml of formic acid solution and added 2g of solid CNBr to start the cleavage reaction. After overnight reaction, the flask was roto-evaporated to dryness. The clear peptide film was resuspended in 40% acetonitrile / 60% H_2_O and mixed for >1h. After centrifugation to remove any insoluble material, the supernatant was lyophilized, resuspended in H_2_O + 0.1% trifluoroacetic acid and run over a C18 reverse phase HPLC column using a gradient of 9-28% acetonitrile. Peak fractions were collected, lyophilized, and analyzed by MALDI-MS to identify the desired peptide product. Dry fractions of peptide were resolublized in DMSO-d6 to a concentration of >5mM.

To measure residual dipolar couplings (RDCs) of α-actinin-1_EF34 or apoCaM bound to IQ peptide, the filamentous bacteriophage Pf1 (Asla Biotech Ltd., Latvia) was used as an orienting medium. Pf1 (12 mg/ml) was added to ^15^N-labeled α-actinin-1_EF34 or apo CaM bound to unlabeled IQ at pH 7.0, to produce weak alignment of the complex.

### Haddock structure determination of α-actinin-1/IQ or apoCaM/IQ

The molecular docking of α-actinin-1 and apoCaM to the Ca_V_1.2 IQ motif (residues 1644 – 1664) was performed using the Haddock d-level 2.2 web server as described (van Zundert et al., 2016). Residual dipolar couplings, chemical shift perturbation, and mutagenesis data were used as structural restraints. For active restraints or ambiguous interaction restraints (AIR), chemical shift perturbation was used, selecting residues whose chemical shift perturbation falls above the average perturbation.

The initial docking calculation used NMR-derived structures of α-actinin-1 or apoCaM as determined in this study, which were each docked with the helical structure of Cav1.2 IQ peptide (residues 1644-1664) (Van Petegem et al., 2005) as input structures for Haddock. A total of 69 and 42 AIR restraints were used for α-actinin-1 and apoCaM, respectively, based on chemical shift perturbation data and 16 active restraints were used for the IQ peptide from chemical shift perturbation. Unambiguous restraints were introduced to define key intermolecular interactions (IQ K1647 - α-actinin-1_EF34 E847; IQ K1647 - α-actinin-1_EF34 E851; IQ K1662 – CaM E88; IQ I1654 - α-actinin-1_EF3/4 F833 and IQ I1654 – apoCaM F90), which were each verified by mutagenesis (Tables 2 and 4).

Initial docking calculations used AIRs based on chemical shift perturbation data and the top 200 structures were selected for simulated annealing and water refinement. The lowest energy structures were then run again, adding unambiguous restraints based on mutagenesis data. Rigid-body docking, simulated annealing, and water refinement were run using the top 200 structures. RDC restraints assigned to α-actinin-1 and apoCaM were then added using the Sani statement, with tensor values Dr and Da calculated using the program PALES (Zweckstetter, 2008). A total of 74 RDC values were used from residues found in regions of regular secondary structure and as deemed reliable by the PALES calculation.

### Fluorescence polarization (FP) assays

Fluorescein-labeled peptides (100 nM; ChinaPeptides, Shanghai, China) were titrated with increasing concentrations of either purified α-actinin-1 or CaM in FP buffer (50 mM HEPES, pH 7.4, 100 mM KCl, 1 mM MgCl_2_, 0.05 mM EGTA, 5 mM nitrilotriacetic acid) and FP determined with a Synergy 2 plate reader (BioTek, Winooski, VT) as described (Patriarchi et al., 2016, Tseng et al., 2017, Zhang et al., 2014). FP was calculated as P = (I_v_ - g*I_h_) / (I_v_ + g*I_h_); I_v_ is vertical and I_h_ are horizontal fluorescence intensity, respectively and g is the correction factor for fluorescein. To obtain binding curves and K_d_ values, data were fitted in GraphPad Prism 5 to the equation Y = B*X / (K_d_ + X); B is maximal FP value that would be reached at saturation as determined by extrapolation of the fitted curve.

### Isothermal titration calorimetry

ITC experiments were performed using a VP-ITC calorimeter (Micro-Cal) at 27°C and data were acquired and processed with MicroCal software as described previously (Wingard et al., 2005). Samples of apoCaM (injectant) and IQ (titrant) were prepared by exchanging each into buffer containing 50 mM HEPES, pH 7.4, 100 mM NaCl, 0.05 mM EGTA and 1 mM MgCl_2_. The IQ peptide in the sample cell (10 μM, 1.5 mL) was titrated with apoCaM (100 μM) using 35 injections of 10 μl each.

### Expression of Ca_V_1.2 and α-actinin-1 in HEK293 cells

HEK293T/17 cells were maintained in DMEM-10 [Dulbecco’s modified Eagle’s medium (Life Technologies) supplemented with 10% fetal bovine serum (FBS, Atlanta Biologicals)] at 37 °C in humidified incubators injected with 5% CO_2_ and 95% air (Tseng et al., 2017). For expression of Ca_V_1.2 HEK293T/17 cells were transfected with rat α_1_1.2 (GenBank ID: M67515.1) cDNA subcloned into pECFP-C1 vector (Tseng et al., 2017) encoding an in frame N-terminally fused eCFP tag and an HA tag in the S5-H5 extracellular loop of domain II (Green et al., 2007), which does not affect channel properties (Altier et al., 2002). The point mutations in plasmids encoding single-residue K1647A, K1647E, F1648A, Y1649A, I1654A, F1658A, and K1662E exchanges in α_1_1.2 were generated via QuikChange II using the above pECFP-C1 rat α_1_1.2 plasmid DNA as template as described (Tseng et al., 2017) and the oligonucleotides described in Suppl. Table 1. For full expression of Ca_V_1.2 cells were also co-transfected with pGWIH-based plasmids encoding the auxiliary subunits rat β_2A_ (Perez-Reyes et al., 1992) and rabbit α_2_δ-1 (Ellis et al., 1988). To assess the contributions of α-actinin-1 on Ca_V_1.2, cells were transfected with pCMV plasmid DNAs encoding WT (Hall et al., 2013) or mutant α-actinin-1. The K847E/K851EE point mutations in α-actinin-1 were produced via QuikChange II mutagenesis as before (Tseng et al., 2017). The forward primer for making the K851E mutation, which was performed first, was 5’p-CCA TGG ACA AAT TGC GCA GAA AGC TGC CAC CCG ACC AGG and reverse primer 5’p-CCT GGT CGG GTG GCA GCT TTC TGC GCA ATT TGT CCA TGG. Forward primer for the K847E mutation was 5’p-GAA CTA CAT TAC CAT GAA CAA ATT GCG CCG CGA GCT GCC ACC C and reverse primer 5’p-GGG TGG CAG CTC GCG GCG CAA TTT GTC CAT GGT AAT GTA GTT C. Transfection of HEK293T/17 cells was accomplished using Ca^2+^ phosphate precipitation (Tseng et al., 2017) for single channel recording and surface labeling or Lipofectamine 2000 for analysis of Q_on_.

#### Cell Surface Biotinylation Assays

Seven hours after transfection, cells were washed once with 150 mM NaCl, 10 mM NaHCO_2_, pH 7.4 (PBS) and cultured for another 15-17 hours in fresh medium. 22-24 h after transfection cells were either harvested, monodispersed (>95% viability) and then labeled with EZ-link-sulfo-NHS-LC-biotin (Thermo Fisher Scientific) in solution essentially as described (Tseng et al., 2017) or directly labeled while adhered to the petri dish. In the latter case, adherent cells on the dish were washed twice with ice cold PBS containing 0.5 mM Mg^2+^ and 1 mM Ca^2+^ and incubated on ice with PBS containing 0.4 mg/mL EZ-link-sulfo-NHS-LC-biotin for 15 min. The labeling reaction was quenched by three five min washes with 40 mM glycine in PBS containing 0.5 mM Mg^2+^ and 1 mM Ca^2+^ on ice. Cells were harvested by scraping in PBS and collected by centrifugation. Cell pellets were lysed in ice-cold RIPA buffer (50 mM Tris-HCl, pH 7.4, 150 mM NaCl, 5 mM EGTA, 10 mM EDTA, 1% NP-40, 0.05% SDS, 0.4% DOC, and 10% glycerol) supplemented with a cocktail of protease inhibitors (1μg/mL leupeptin (Merck Millipore), 2μg/mL aprotinin (Merck Millipore), 1 μg/mL pepstatin A (Merck Millipore), and 34 μg/mL phenylmethanesulfonyl fluoride (PMSF; Sigma)). The solubilized material was cleared of insoluble debris by centrifugation at 200,000xg for 30 min at 4°C. Biotinylated constituents in 600 μg of cell protein lysate were affinity purified by incubation with 30 μL of NeutrAvidin conjugated Sepharose beads (Thermo Fisher Scientific) for 2 h at 4°C. Bead-bound material was collected by centrifugation and washed three times with ice-cold buffer, and immobilized proteins were extracted in SDS sample buffer. Proteins were fractionated by SDS-PAGE in 7.5% acrylamide gels and transferred onto polyvinyldene difluoride (PVDF; BioRad) membranes. PVDF membranes were incubated in blocking buffer (BB) consisting of 150 mM NaCl, 10 mM Tris-Cl, pH 7.4 (TBS) with 0.10% Tween (TBST) and 2% bovine serum albumin (BSA; RPI Corp.) for 1 h at RT before incubation with primary antibodies in BB for 3 h at RT. α_1_1.2 was detected by antibodies against the HA tag and, for confirmation, the intracellular loop II/III FP1 epitope (Buonarati et al., 2017) and the CNC2 epitope near the C-terminus of α_1_1.2 (Buonarati et al., 2017). Probing with antibodies against the cytosolic protein vinculin (Cell Signaling Technologies) and α-tubulin (Santa Cruz Biotechnology) were used to correct for variation in protein content in lysates and as negative control for surface biotinylation. Membranes were washed for 40 min with at least five exchanges of TBST, incubated with horseradish peroxidase-conjugated secondary goat anti-mouse antibodies (HA, α-tubulin; Jackson) or mouse anti-rabbit antibodies (FP1, CNC2, vinculin; Jackson) for 1 h at RT, and washed again with TBST with at least five exchanges for 1.5 h. Immunosignals were detected using the horseradish peroxidase substrates Luminata Classico or Crescendo (Merck Millipore) or Femto (Thermo Fisher Scientific) by X-ray film (Denville Scientific Inc.). Multiple exposures over increasing time periods were taken to ensure that all signals were in the linear range (Davare and Hell, 2003, Hall et al., 2006). Films were scanned and assessed via image J to determine signal intensity for each band. Background signals in individual lanes were subtracted from the band signal before further analysis. To correct differences in immunosignal strengths due to potential differences during immunoblotting and film exposures between experiments, the individual immunosignals for each α_1_1.2 pull-down sample were divided by the sum of all immunosignals from one blot to obtain the relative signal fraction for each band (Degasperi et al., 2014). The mean of the WT control signals from all experiments was then calculated and all fractional values from all samples (WT and mutants) divided by this value (the WT mean transforms to 1 with this algorithm) (Degasperi et al., 2014). All values were then converted to percent, with the mean of WT control equaling 100%. The data were statistically analyzed applying either a student’s T-test (two sample comparison) or ANOVA with Bonferroni’s post hoc test as before (Tseng et al., 2017).

### Cell-attached Patch Clamp Recording

Cell-attached patch clamp recordings were performed as before (Davare et al., 2001, Patriarchi et al., 2016, Qian et al., 2017) on an Olympus IX70 inverted microscope at room temperature (22°C). Recordings were obtained with an Axopatch 200B amplifier and data were sampled at 10 kHz with a low-pass filter at 2 kHz (3 dB, four pole Bessel) and digitalized with a Digidata 1440 digitizer. Recording electrodes were fabricated from borosilicate capillary glass (0.86 OD) with a Flaming micropipette puller (Model P-97, Sutter Instruments) and polished (polisher from World Precision Instruments; 3.5 - 6.5 MΩ resistance). The extracellular solution contained (in mM) 145 KCl, 10 NaCl, and 10 HEPES, pH 7.4 (NaOH). The high K^+^ concentration was used for optimal control of the transmembrane potential under the patch during depolarizations to 0 mV. The pipette solution contained (in mM) 20 tetraethylammonium chloride (TEA-Cl), 110 BaCl_2_ (as charge carrier), and 10 HEPES, pH 7.3 (TEA-OH). Cells were depolarized from a holding potential of −80 mV to 0 mV every 5 seconds. Event lists were translated from raw Ba^2+^ currents after leak and capacity transients were digitally subtracted. Data were analyzed based on the half-hight criterium (Sachs et al., 1982) with the single channel software provided by pClamp 10. The number of channels (*k*) in the patch were estimated based on the observed simultaneous and staged openings) over several minutes at the depolarizing test potential (Bartels et al., 2018, Herzig et al., 2007). The *k*-value is then determined by the observed maximum current amplitude divided by the unitary current amplitude. Data with more than >3 channels (*k* > 3) in the patch were not considered for statistical analysis in order to prevent overinterpretation of channel open probability. For statistical analysis single channel parameters were corrected by the channel number as previously (Bartels et al., 2009, Schroder et al., 1998). For a sufficient statistical analysis 50-100 Ba^2+^ current traces were recorded on average for each cell for each experimental condition.

### Whole-cell patch-clamp recordings

Whole-cell patch clamp recordings were performed as before (Bartels et al., 2018) on an Olympus IX70 inverted microscope at room temperature (22°C). Macroscopic Ba^2+^ currents (I_Ba_) of Ca_v_1.2 were recorded in external solution containing (in mM) 75 CsCl, 40 TEA-Cl, 20 BaCl_2_, 1 MgCl_2_, 10 HEPES and 10 Glucose with a pH adjusted to 7.2 (TEA-OH) and an osmolarity of 300-310 (sucrose). The internal pipette solution contained (in mM) 110 CsCl, 30 TEA-Cl, 1 MgCl_2_, 4 Mg-ATP, and 10 HEPES, pH 7.2 (CsOH), mOsm 290-300 (sucrose). Pipette resistance was usually between 1.7-2.5 MΩ. The series resistance and the cell capacitance were taken from an Axopatch 200B Amplifier (Molecular Device) and compensated not more than <40% in order to prevent current oscillation. On-gating currents (Q_on_) were sampled at 50 kHz and lowpass-filtered at 5 kHz and further quantified through current integration over the first 2-3 ms of the beginning of the test pulse. Cells were clamped at a holding potential of −80 mV and depolarized by a 20 ms test pulse of a serious of activating potentials starting from −60 mV to +80 mV to determine the reversal potential (E_rev_). Tail currents (I_tail_) were then measured after repolarization to −50 mV for 10 ms. Recorded data were leak and capacity corrected with an online *P/4* protocol. The liquid junction potential was not considered for correction in the experiment. Data acquisition and analysis was obtained with pClamp 10. Curve fitting was performed by using GraphPad Prism VIII software (San Diego).

## Acknowledgements

We thank Dr. Elza Kuzmenkina (University of Cologne, Germany) for providing the coding of the algorithm for calculating Assembly Averages.

This work was supported by NIH grants T32 GM113770 (AMC), T32 GM099608 (PBH), R01 HL098200 (MFN), R01 HL121059 (MFN), R01 EY012347 (JBA), R01MH097887 (JWH), R01 AG017502 (JWH), R01 NS078792 (JWH), and the American Heart Association Predoctoral Fellowship AHA 14PRE19900021 (PBH).

## Author Contribution

MT, DEA, MN-C, PB, MFN, MCH, JBA, and JWH designed experiments, MT, DEA, MN-C, PB, AMC, PBH, MFN, and MCH performed experiments, MT, DEA, MN-C, PB, AMC, PBH, VY-Y, DMB, MFN, MCH, JBA, and JWH analyzed data, and MT, DEA, MN-C, PB, VY-Y, DMB, MFN, MCH, JBA, and JWH wrote the manuscript.

## Data Availability

The NMR assignments have been deposited in the BMRB (accession number 25902). The atomic coordinates have been deposited into the Protein Databank (6COA and 6CTB).

Supplemental Material is available in the Online version of this article.

## SUPPLEMENTAL MATERIAL

**Supplemental Table 1.**
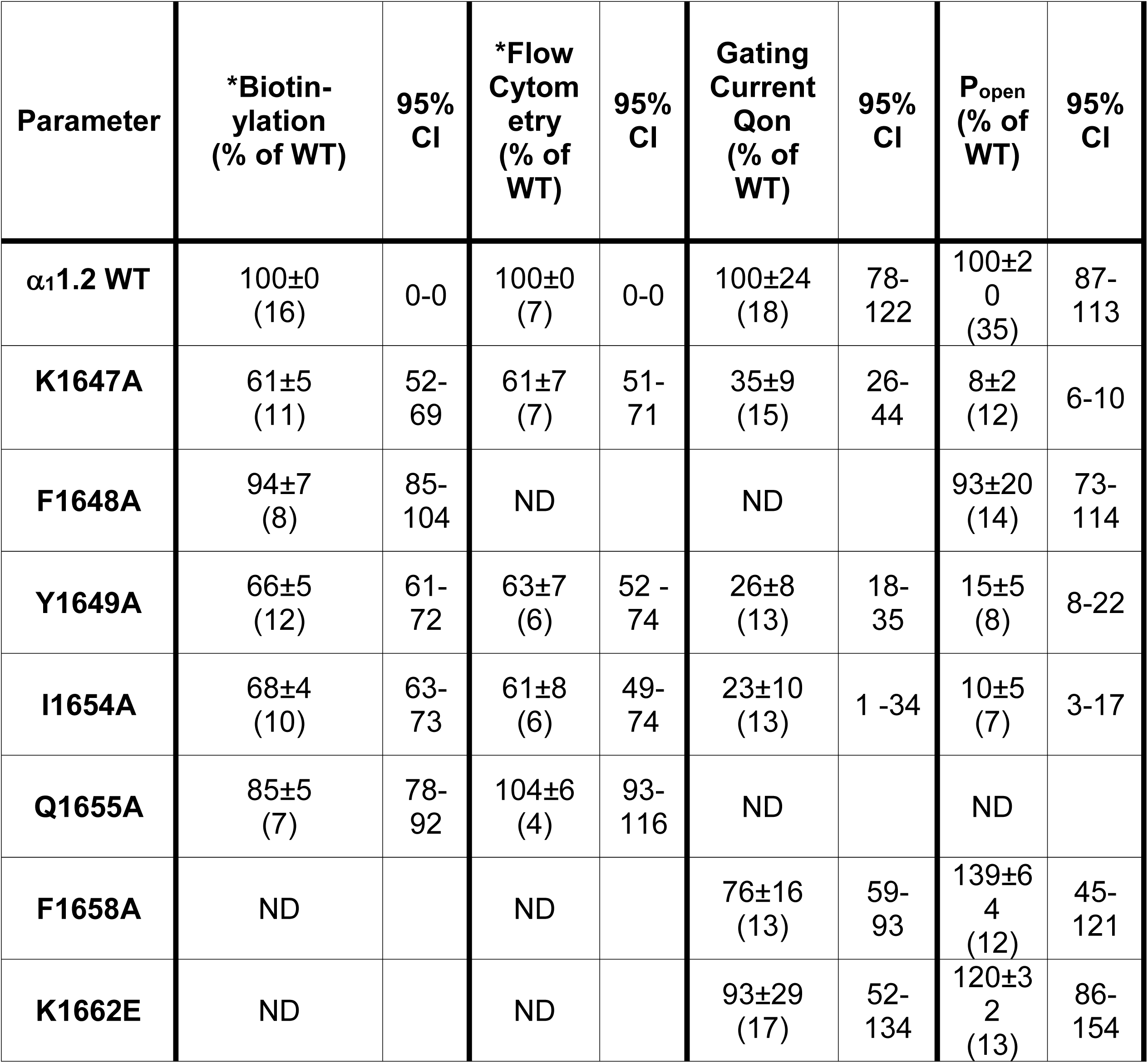
95% Confidence intervals (CI) for surface labelling, Qon, and Po for WT and IQ mutant Ca_V_1.2. Given are means±SEM and 95% CIs for experimental values. The number of experiments is shown in parenthesis. *The biotinylation and flow cytometry data are based on data originally published by Tseng et al (2017).

**Supplemental Table 2.**
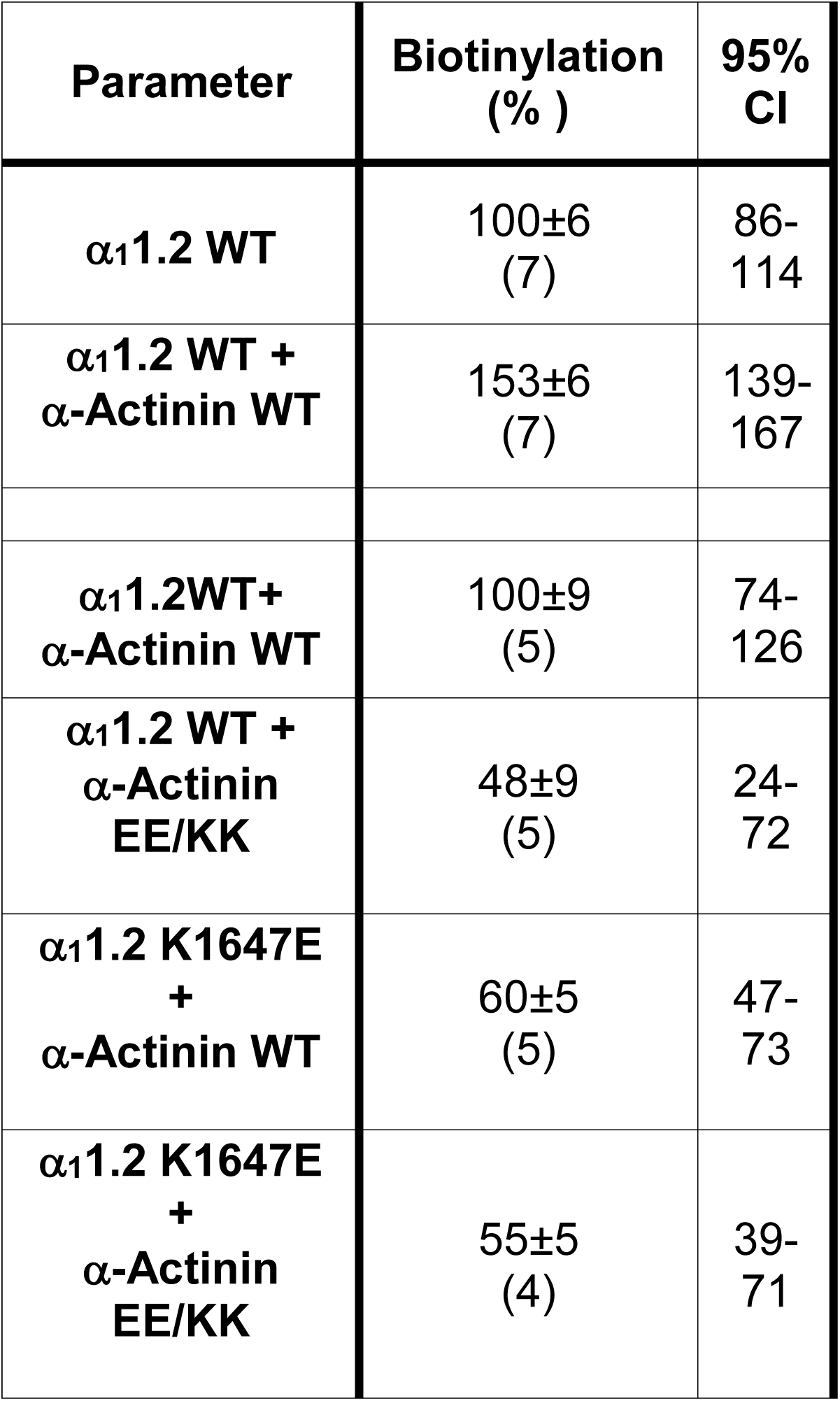
95% Confidence intervals (CI) for surface biotinylation for Ca_V_1.2 / α-actinin-1 charge inversion experiments. Given are means±SEM and 95% CIs (based on original data from this manuscript). The number of experiments is shown in parenthesis.

**Supplemental Table 3.** 95% Confidence intervals (CI) for Po for Ca_V_1.2 / α-actinin-1 charge inversion experiments. Given are means±SEM and 95% CIs (based on original data from this manuscript). The number of experiments is shown in parenthesis.

| Parameter | P<sub>open</sub> (%) | 95% CI |
| --- | --- | --- |
| $\alpha_1$1.2 WT | 100±20<br>(35) | 87-113 |
| $\alpha_1$1.2 WT +<br>α-Actinin WT | 233±59<br>(23) | 186 -<br>280 |
| $\alpha_1$1.2 WT+<br>α-Actinin<br>EE/KK | 55±22<br>(13) | 32-<br>78 |
| $\alpha_1$1.2 K1647E<br>+<br>α-Actinin WT | 100±18<br>(30) | 87-113 |
| $\alpha_1$1.2K1647E<br>+<br>α-Actinin<br>EE/KK | 147±28<br>(16) | 120-174 |

**Supplemental Figure 1. Amino acid sequence alignment of the membrane proximal portion of the C-terminal tail of L-type Ca^2+^ channels.**

Bold refers to reference sequence (rat α_1_1.2), turquoise to divergency in amino acid sequence, red to residues K1647 and Y1649, which are important for α-actinin binding only, green to I1654, which is important for both, α-actinin and apoCaM binding, and yellow to F1658 and K1662, which are important for apoCaM binding only. Amino acid sequences were derived from species according to the numerical labeling^1-13^. Data were extracted and compiled from uswest.ensembl.org.

**Supplemental Figure 2. Mapping the Ca_V_1.2 IQ binding site in α-actinin and apoCaM.**

a, Overlay of ^15^N-^1^H HSQC spectra of ^15^N-labeled α-actinin-1 CH1/CH2 domain (residues 19-192) by itself (black peaks) and after addition of saturating, unlabeled IQ peptide (red peaks). b, Overlay of ^15^N-^1^H HSQC spectra of ^15^N-labeled α-actinin-1 EF-hand domain (residues 750-892) by itself (black peaks) and after addition of saturating, unlabeled IQ peptide (red peaks). c, Overlay of ^15^N-^1^H HSQC spectra of ^15^N-labeled α-actinin-1 C-lobe (α-actinin-1 EF34; residues 822-892) by itself (black peaks) and after addition of saturating, unlabeled IQ peptide (red peaks).

d, Overlay of ^15^N-^1^H HSQC spectra of ^15^N-labeled apoCaM by itself (black peaks) and after addition of saturating, unlabeled IQ peptide (red peaks).

**Supplemental Figure 3. NMR structural analysis using residual dipolar couplings (RDCs).**

a,b, ^15^N-^1^H HSQC-IPAP spectra (Tjandra and Bax, 1997) of ^15^N-labeled α-actinin-1_EF34 (A) or apoCaM (B) in the presence of saturating unlabeled IQ peptide. Representative and measured RDC values are marked above selected peaks. Samples used in HSQC-IPAP experiments were prepared by adding 5 mg of ^15^N-labeled protein to 0.5 mL of NMR buffer containing 12 mg/mL of filamentous bacteriophage Pf1. HSQC-IPAP spectra were recorded in the presence of Pf1 (a,b) and absence of Pf1 (not shown). Residual dipolar couplings (RDCs) were measured (in Hz) as the difference in splitting for the ^15^N-[^1^H] doublet components relative to the isotropic ^1^J_NH_ coupling. RDCs measured for 34 residues (α-actinin-1_EF34/IQ) and 36 residues (apoCaM/IQ) served as orientational structural restraints applied during the refinement phase of the structure calculation (Schwieters et al., 2003).

(C-D) Plots showing the correlation between observed versus calculated backbone RDCs predicted from the final NMR derived structures for α-actinin-1_EF34/IQ and apoCaM/IQ, respectively. The correlation coefficient (r^2^) equals 0.99 and Q is 0.095 and 0.089 for α-actinin-1_EF34/IQ and apoCaM/IQ, respectively.

**Supplemental Figure 4. Isothermal titration calorimetry of apoCaM added to IQ peptide.**

a, Change of heat resulting from incremental addition of apoCaM (100 μM stock solution) into IQ peptide (10 μM) during ITC titration at 27°C in 50 mM HEPES (pH 7.4), 100 mM KCl, 0.05 mM EGTA and 1 mM MgCl_2_.

b, A binding isotherm of apoCaM binding to IQ peptide was derived from the integrated heat at each injection after subtracting a blank titration (to remove heat of dilution). The binding isotherm for apoCaM binding to IQ exhibited no detectable heat signal above the noise level, consistent with low fractional binding under ITC conditions and/or low enthalpy caused by the relatively weak binding affinity (K_d_ = 10 μM, see Fig. 2e and Table 4).

c, Fluorescence polarization for binding of apoCaM (0.015 to 256 μM) to fluorescein-labeled WT IQ peptide (1.0 μM) in the presence of 100 mM KCl (black squares) versus zero KCl (red dots) at room temperature. The buffer was the same as in (a) except that KCl was excluded in the zero KCl sample. The data were fit to a one-site model (dotted lines) with K_d_ = 10 μM in the presence of 100 mM KCl and K_d_ = 2.6 μM in the absence of KCl.

**Supplemental Figure 5. Representative single channel recordings and assembly averages for Ca_V_1.2 WT, K1647A, F1648A, Y1649A, I1654A, F1658A, and K1662E.**

HEK293 cells were transfected with α_1_1.2, α_2_δ-1, and β_2A_ before cell attached patch recording. Holding potential (HP) was −80 mV and test potential (TP) 0 mV. Shown are 20 consecutive sweeps from representative experiments (top) and mean assembly averages for all experiments for Ca_V_1.2 WT and each mutant.

a, Ca_V_1.2 WT versus K1647A, F1648A, Y1649A

b, Ca_V_1.2 WT versus I1654A, F1658A, and K1662E.

Ca_V_1.2 WT in and K1647A are replicated from Fig. 3 a,b for comparison with the other Ca_V_1.2 mutants.

**Supplemental Figure 6. Effects of Ca_V_1.2 IQ mutants on single channel properties.**

HEK293 cells were transfected with α_1_1.2, α_2_δ-1, and β_2A_ before cell attached patch recording. Shown are dot plots and means ± SEM (whiskers) for availability (a), NPo (b), Po (c), the mean of the currents calculated from the ensemble averages of each experiment (d), and single channel conductances i (e) for Ca_V_1.2 WT, K1647A, F1648A, Y1649A, I1654A, F1658A, and K1662E

(**p<0.01, ***p<0.001 compared to WT; one-way ANOVA with Bonferroni post-hoc test (a) or Welch ANOVA with Tamhane T2 test (b-e).

**Supplemental Figure 7. Effects of overexpression of α-actinin-1 on single channel properties of WT Ca_V_1.2.**

HEK293 cells were transfected with WT α_1_1.2, α_2_δ-1, and β_2A_ plus, if indicated, α-actinin-1 WT or E847K/E851K before cell attached patch recording.

Shown are dot plots and means ± SEM (whiskers) for availability (a), NPo (b), Po (c), the mean of the currents calculated from the ensemble averages of each experiment (d), and single channel conductances i (e; *p<0.05, **p<0.01, ***p<0.001; unpaired t-test).

**Supplemental Figure 8. α-Actinin-1 E847K/E851K rescues open probability for Ca_V_1.2 K1647E.**

HEK293 cells were transfected with K1647E α_1_1.2, α_2_δ-1, and β_2A_ plus α-actinin-1 WT or E847K/E851K before cell attached patch recording.

Shown are dot plots and means ± SEM (whiskers) for availability (a), NPo (b), Po (c), the mean of the currents calculated from the ensemble averages of each experiment (d), and single channel conductances i (e; **p<0.01; unpaired t-test).

**Supplemental Figure 9. Ca_V_1.2 mutations that affect α-actinin-1 impair gating charge movement and its coupling to channel opening.**

HEK293 cells were transfected with α_1_1.2, α_2_δ-1, and β_2A_ before whole cell patch recording in 20 mM Ba^2+^.

a, Dot plots and means ± SEM (whiskers) of Q_on_ determined at reversal potential (see Fig. 7 for details; *p<0.05; **p<0.01; one-way ANOVA with Bonferroni post-hoc test).

b, I-V curves. Currents were recorded upon depolarization from a holding potential of −80 mV to increasingly more positive potentials (insert on left). Shown are peak currents. Dashed lines indicate SEM. The WT Cav1.2 curve is reproduced in all graphs for the Ca_V_1.2 mutants.

c, G-V curves. Tail currents (I_Tail_) were recorded upon repolarization to −50 mV following depolarization from a holding potential of −80 mV to increasingly more positive potentials. Shown are fitted curves in tope panels and dot blots in bottom panels.

**Supplemental Figure 10. IQ peptide binds with opposite polarity to α-actinin-1 versus apoCaM.**

a, NMR structure of α-actinin-1 (cyan) bound to Ca_V_1.2 IQ motif (red). b, NMR structure of apoCaM (cyan) bound to Ca_V_1.2 IQ motif (red).

c, NMR structure apoCaM (cyan) bound to the Na_V_1.2 IQ motif (red). PDB accession number is 2KXW (Feldkamp et al., 2011). Non-conserved intermolecular contacts between hydrophobic side chain atoms are colored yellow. The β-methyl side chain atoms of Na_V_1.2 residues A1909 and A1915 are 2.5Å away from side chain methyl atoms of apoCaM residues M109 and L85, respectively. The aromatic side chain atoms of Na_V_1.2 residue Y1919 are 3.3 Å away from aromatic side chain atoms of apoCaM residue F141.

a-c, Black arrows depict directionality of the IQ helix.

d, Amino acid sequence alignment of IQ motifs from Na_V_1.2, Na_V_1.5, and Ca_V_1.2. Residues that are conserved between Na_V_ and Ca_V_ are shown in boldface font. Lysine residues in Ca_V_1.2 that form intermolecular salt bridges in the Ca_V_1.2 IQ peptide complexes with α-actinin-1_EF3/4 and apoCaM and are not conserved between the Ca^2+^ and Na^2+^ channels are colored red.

**Supplemental Figure 11. Structural modeling addressing the function of Y1649 in the α_1_1.2 IQ motif and how the E851K mutation of α-actinin affects its binding to the IQ motif**

a, Y1649 stabilizes the α-helical conformation of the α_1_1.2 IQ motif and thereby α-actinin binding. Shown is a cartoon representation of α_1_1.2 IQ motif α-helix (orange) and α-actinin-1 EF3-EF4 (blue). Sidechains of key residues are depicted as stick representation and labeled. Y1649 is in close proximity to L1653, which is about one α-helical turn downstream of Y1649. Stabilization of this α helix ensures correct positioning of I1654, which is critical for binding to α-actinin. This figure was created using UCSF Chimera (Pettersen et al., 2004).

b-d, The lysine residue in the E851K mutation of α-actinin destabilizes the canonical EF hand structure. Conformers of the lysine residues when used to substitute E851 by itself or together with E847 in α-Actinin-1 and of the glutamate residue when used to substitute K1647 in the IQ motif were calculated for the structure of the complex between the IQ motif of α_1_1.2 and EF3_EF4 of α-Actinin-1 using the UCSF Chimera rotamer tool (Pettersen et al., 2004). The α_1_1.2 IQ α-helix is shown in orange and the α-actinin-1 EF3-EF4 in blue. Sidechains of key residues are depicted as stick representation and labeled. The lysine rotamers (thin sticks) are overlaid on top of the original E851 and E847 residues and the glutamate rotamers on top of the original K1647 residue.

b, Simulation of rotamers for the E851K mutation in α-actinin-1 to show possible conformations of the lysine sidechain.

c, Simulation of rotamers for the E851K mutation in α-actinin-1 and the K1647E mutation in α_1_1.2 to show possible conformations of the lysine and glutamate sidechains.

d, Simulation of rotamers for the E847K and E851K mutations in α-actinin-1 and the K1647E mutation in α_1_1.2 to show possible conformations of the lysine and glutamate sidechains.

