## Supplemental Figure 1 for "α-Actinin-1 promotes activity of the L-type Ca^2+^ Channel Ca_V_1.2"

```
Species                 --------Pore---------------   ----------------------C-terminus------------------------------------

                            IVS6                           EF-hand:++   ++    ++ + ++   --   ++   ++    +
Human        Cav1.1  AVFYFISFYMLCAFLIINLFVAVIMDNFDYLTRDWSILGPHHLDEFKAIWAEYDPEAKGRIKHLDVVTLLRRIQPPLGFGKFCPHRVACKRLVGMNMP
Human        Cav1.2  AVFYFISFYMLCAFLIINLFVAVIMDNFDYLTRDWSILGPHHLDEFKRIWAEYDPEAKGRIKHLDVVTLLRRIQPPLGFGKLCPHRVACKRLVSMNMP
Human        Cav1.3  AIVYFISFYMLCAFLIINLFVAVIMDNFDYLTRDWSILGPHHLDEFKRIWSEYDPEAKGRIKHLDVVTLLRRIQPPLGFGKLCPHRVACKRLVAMNMP
Human        Cav1.4  AIAYFISFFMLCAFLIINLFVAVIMDNFDYLTRDWSILGPHHLDEFKRIWSEYDPGAKGRIKHLDVVALLRRIQPPLGFGKLCPHRVACKRLVAMNMP
Monkey       Cav1.2  AVFYFISFYMLCAFLIINLFVAVIMDNFDYLTRDWSILGPHHLDEFKRIWAEYDPEAKGRIKHLDVVTLLRRIQPPLGFGKLCPHRVACKRLVSMNMP
Cow          Cav1.2  AVFYFISFYMLCAFLIINLFVAVIMDNFDYLTRDWSILGPHHLDEFKRIWAEYDPEAKGRIKHLDVVTLLRRIQPPLGFGKLCPHRVACKRLVSMNMP
Rabbit       Cav1.2  AVFYFISFYMLCAFLIINLFVAVIMDNFDYLTRDWSILGPHHLDEFKRIWAEYDPEAKGRIKHLDVVTLLRRIQPPLGFGKLCPHRVACKRLVSMNMP
Rat          Cav1.2  AVFYFISFYMLCAFLIINLFVAVIMDNFDYLTRDWSILGPHHLDEFKRIWAEYDPEAKGRIKHLDVVTLLRRIQPPLGFGKLCPHRVACKRLVSMNMP
Mole Rat     Cav1.2  AVFYFISFYMLCAFLIINLFVAVIMDNFDYLTRDWSILGPHHLDEFKRIWAEYDPEAKGRIKHLDVVTLLRRIQPPLGFGKLCPHRVACKRLVSMNMP
Mouse        Cav1.2  AVFYFISFYMLCAFLIINLFVAVIMDNFDYLTRDWSILGPHHLDEFKRIWAEYDPEAKGRIKHLDVVTLLRRIQPPLGFGKLCPHRVACKRLVSMNMP
Zebrafish    Cav1.2  AIFYFVSFYMLCAFLIINLFVAVIMDNFDYLTRDWSILGPHHLDEFKRIWAEYDPEAKGRIKHLDVVTLLRRIQPPLGFGKLCPHRVACKRLVSMNMP
Fly          Cav1.2  ---YFISFYVLCSFLIINLFVAVIMDNFDYLTRDWSILGPHHLDEFIRLWSEYDPDAKGRIKHLDVVTLLRKISPPLGFGKLCPHRMACKRLVSMNMP
Sea urchin           AYVYFISFYSICSFLIINLFVAVIMDNFDYLTRDWSILGPHHLDEFVRQWSEFDPDATGRIKHLDVVTLLRSISPPLGFGKLCPHRIACKRLVTMNMP
Snail                AIPYFISFLLLVAFFVLNMFVGVVVDNFDYLTRDWSILGPHHLDEFVRLWSEYDPEAKGRIHYTDMYEMLRNMEPPVGFGKKCP-YKLAQRLVSMNMP
C. elegans           AIVYFISFFMLCSFLVINLFVAVIMDNFDYLTRDWSILGPHHLEEFVRLWSEYDPDAKGRIKHLDVVTLLRKISPPLGFGKLCPHRLACKRLVSMNMP
Jellyfish            AYLYFMSFYMICSFLIINLFVAVIMDNFDYLTRDWSILGAHHLEEYVRIWAEYDPEASGRMKHVDIVSMLKRIEPPLGFGKCCPHREACKRLVSMNMM

                     ------------------------------------C-terminus--------------------------------------------------
                                          Pre-IQ Domain                                  IQ DOMAIN

Human        Cav1.1  LNSDGTVTFNATLFALVRTALKIKTEGNFEQANEELRAIIKKIWKRTSMKLLDQVIPPIG-DDEVTVGKFYATFLIQEHFRKFMKRQEEYYGYRP
Human        Cav1.2  LNSDGTVMFNATLFALVRTALRIKTEGNLEQANEELRAIIKKIWKRTSMKLLDQVVPPAG-DDEVTVGKFYATFLIQEYFRKFKKRKEQGLVGKP
Human        Cav1.3  LNSDGTVMFNATLFALVRTALKIKTEGNLEQANEELRAVIKKIWKRTSMKLLDQVVPPAG-DDEVTVGKFYATFLIQDYFRKFKKRKEQGLVGKY
Human        Cav1.4  LNSDGTVTFNATLFALVRTSLKIKTEGNLEQANQELRIVIKKIWKRMKQKLLDEVIPPPD-EEEVTVGKFYATFLIQDYFRKFRRRKEKGLLGND
Monkey       Cav1.2  LNSDGTVMFNATLFALVRTALRIKTEGNLEQANEELRAIIKKIWKRTSMKLLDQVVPPAG-DDEVTVGKFYATFLIQEYFRKFKKRKEQGLVGKP
Cow          Cav1.2  LNSDGTVMFNATLFALVRTALRIKTEGNLEQANEELRAIIKKIWKRTSMKLLDQVVPPAG-DDEVTVGKFYATFLIQEYFRKFKKRKEQGLVGKP
Rabbit       Cav1.2  LNSDGTVMFNATLFALVRTALRIKTEGNLEQANEELRAIIKKIWKRTSMKLLDQVVPPAG-DDEVTVGKFYATFLIQEYFRKFKKRKEQGLVGKP
Rat          Cav1.2  LNSDGTVMFNATLFALVRTALRIKTEGNLEQANEELRAIIKKIWKRTSMKLLDQVVPPAG-DDEVTVGKFYATFLIQEYFRKFKKRKEQGLVGKP
Mole Rat     Cav1.2  LNSDGTVMFNATLFALVRTALRIKTEGNLEQANEELRAIIKKIWKRTSMKLLDQVVPPAG-DDEVTVGKFYATFLIQEYFRKFKKRKEQGLVGKP
Mouse        Cav1.2  LNSDGTVMFNATLFALVRTALRIKTEGNLEQANEELRAIIKKIWKRTSMKLLDQVVPPAG-DDEVTVGKFYATFLIQDYFRRFKKRKEQGLVGKP
Zebrafish    Cav1.2  LNSDGTVMFNATLFALVRTALRIKTEGNLEQANEELRAIVKKIWKRTSMKLLDQVVPPAG-DDEVTVGKFYATFLIQEYFRKFKKRKEQEGKEGH
Fly          Cav1.2  LNSDGTVLFNATLFAVVRTSLSIKTDGNIDDANSELRATIKQIWKRTNPKLLDQVVPPPGNDDEVTVGKFYATYLIQDYFRRFKKRKEQ------
Sea urchin           LNSDGTVMFNATLFALIRTSLKIKTEGNIDQCNEELRAVIKKIWKRTSTKLLDQVAPPAGADDDVTVGKFYATFLIQDYFRRFKKRKQEGL----
Snail                LNSDGTVMFNATLFALVRTSLKIKTEGNIDTANEELRTVIKKIWKRTSPKLLDQVVPPAG-DDDVTVGKFYATFLIQDYFRRFKKRKEQ------
Worm                 LNSDGTVCFNATLFALVRTNLKIYTEGNIDEANEQLRSAIKRIWKRTHKDLLDEVVPPAGKEDDVTVGKFYATFLIQDYFRRFKKRKE-------
Jellyfish            MNNDGTVDFHATLFALVRTSLNIKKPDAILHANNELRGILKHLWPRTNENLFDKLIPPPDYAEGITVGKFYATFLIQEYFRKFKKKRQEERKKK
```

[1]*Homo sapiens*,[2]*Pan troglodytes*,[3]*Bos taurus*, [4]*Oryctolagus cuniculus*, [5]*Rattus norvegicus*, [6]*Heterocephalus glaber*, [7]*Mus musculus*, [8]*Danio rerio*, [9]*Drosophila melanogaster*, [10]*Strongylocentrus purpuratus*, [11]*Lymnea stagnalis*, [12]*Caenorhabditis elegans* [13]*Cyanea lamarkii*

Color code of residues: **Different from Cav1.2** | **Important for α-actinin binding** | **Important for α-actinin and CaM binding** | **Important for apoCaM binding**
