## Supplementary figures and images for "α-Actinin-1 promotes activity of the L-type Ca^2+^ Channel Ca_V_1.2"

### Supplemental Figure 2

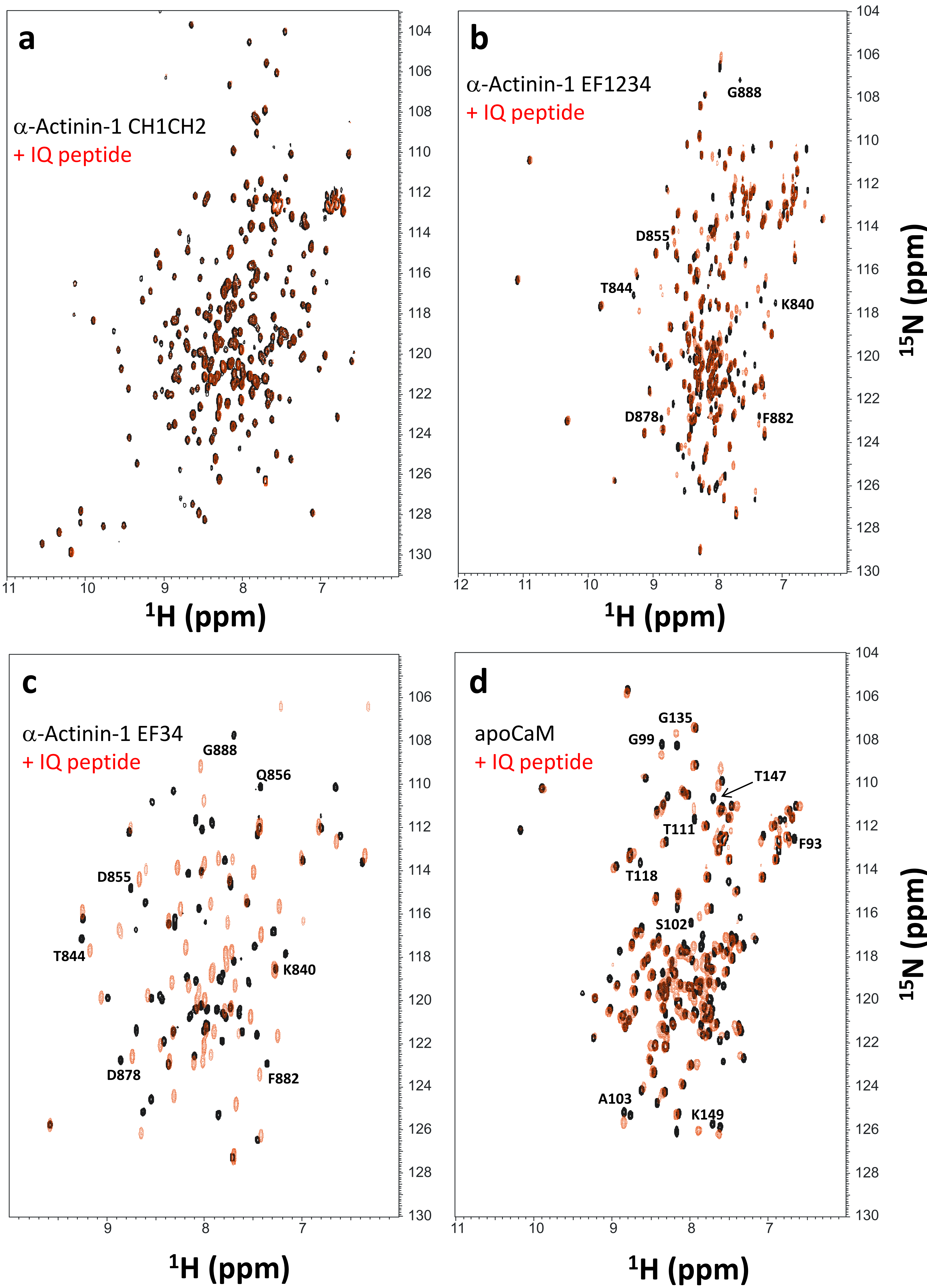

### Supplemental Figure 3

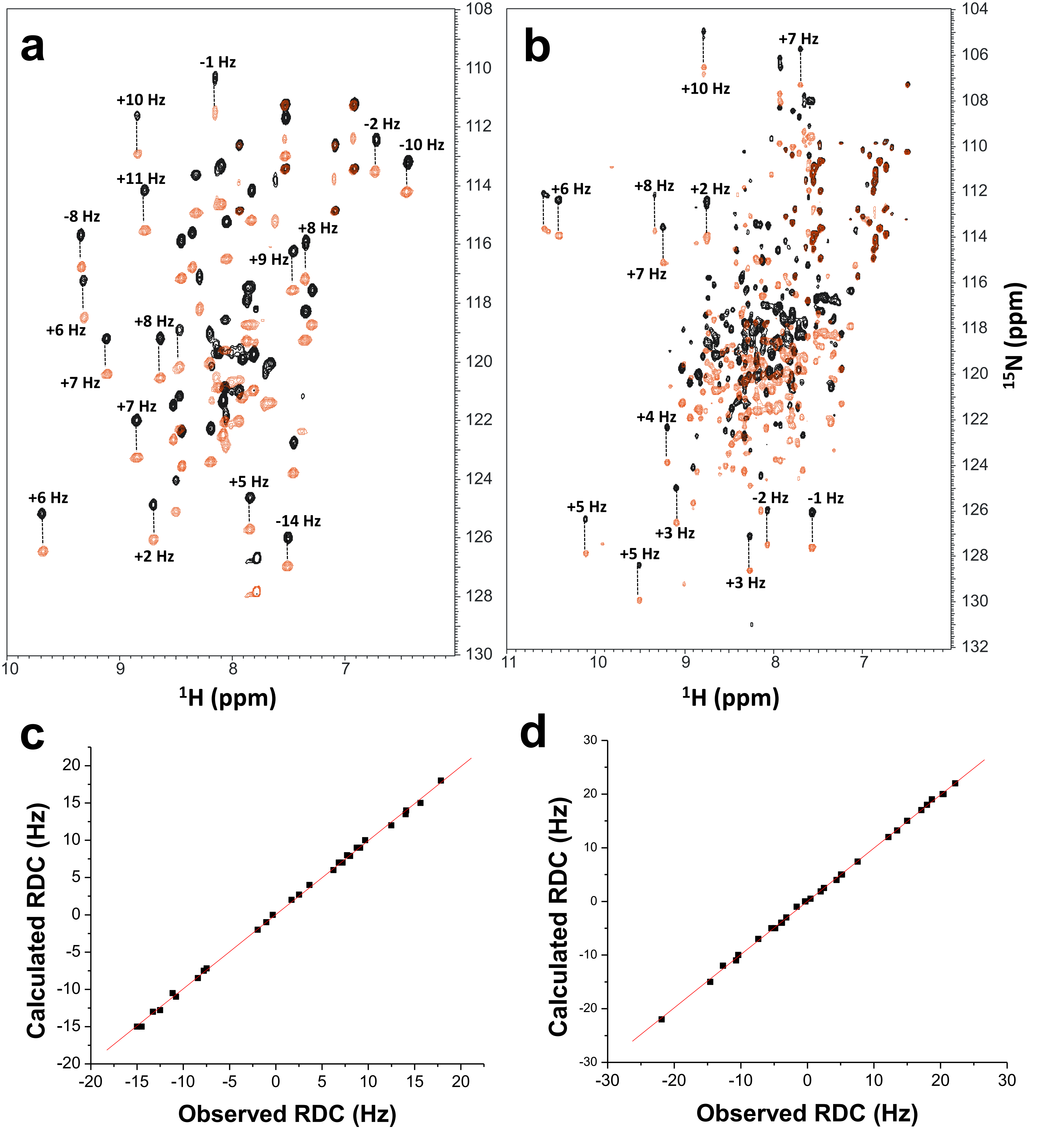

### Supplemental Figure 4

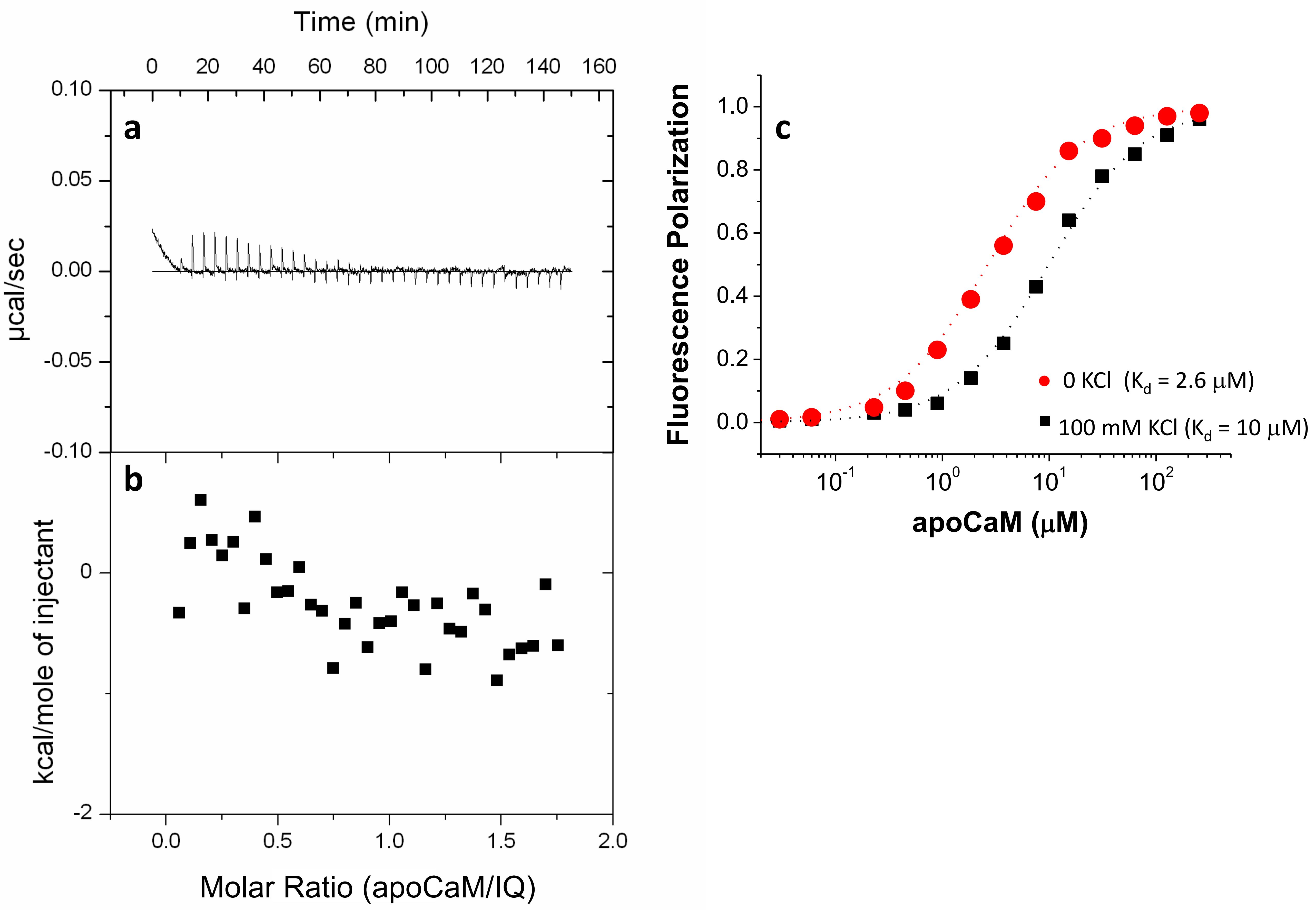

### Supplemental Figure 5a

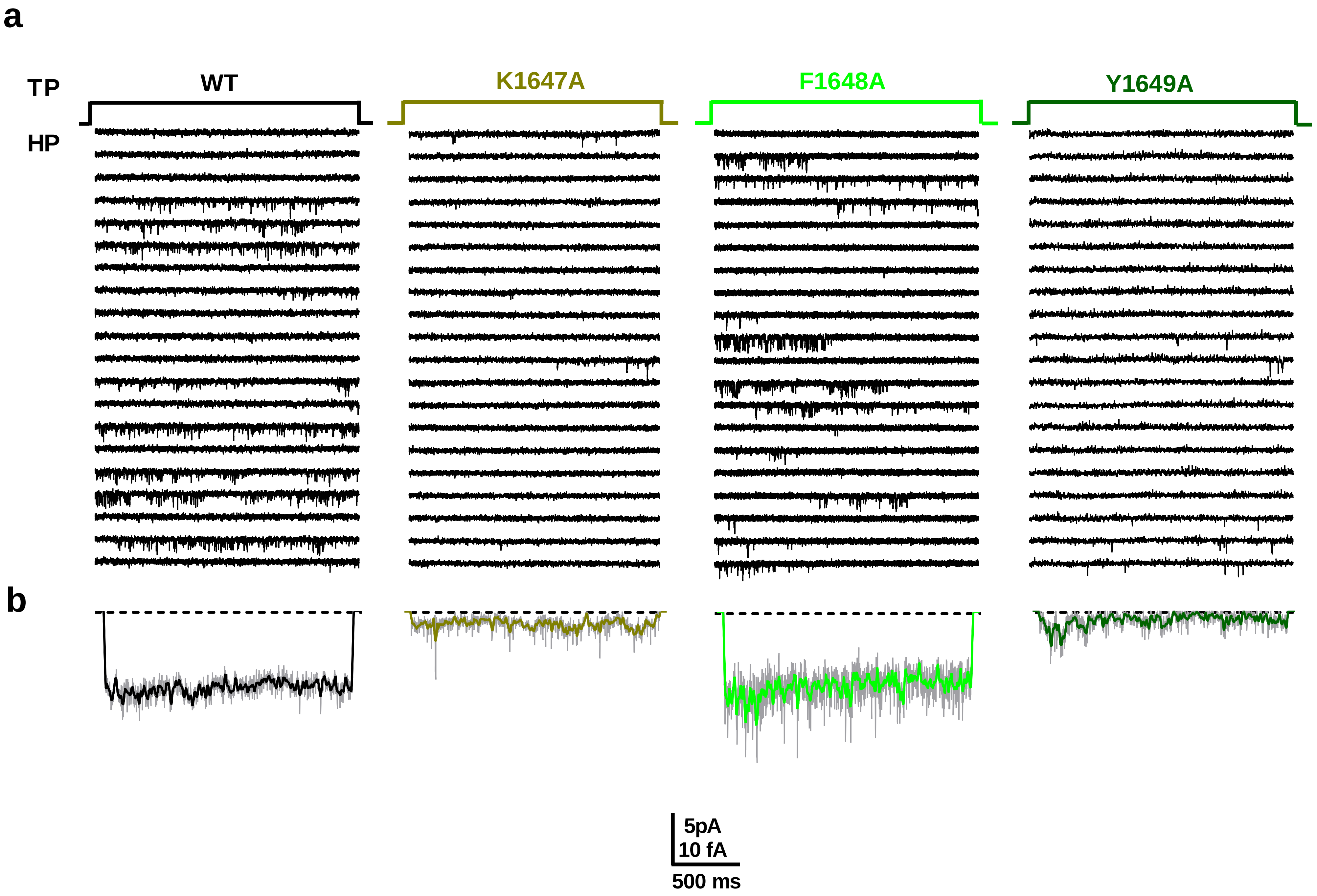

### Supplemental Figure 5b

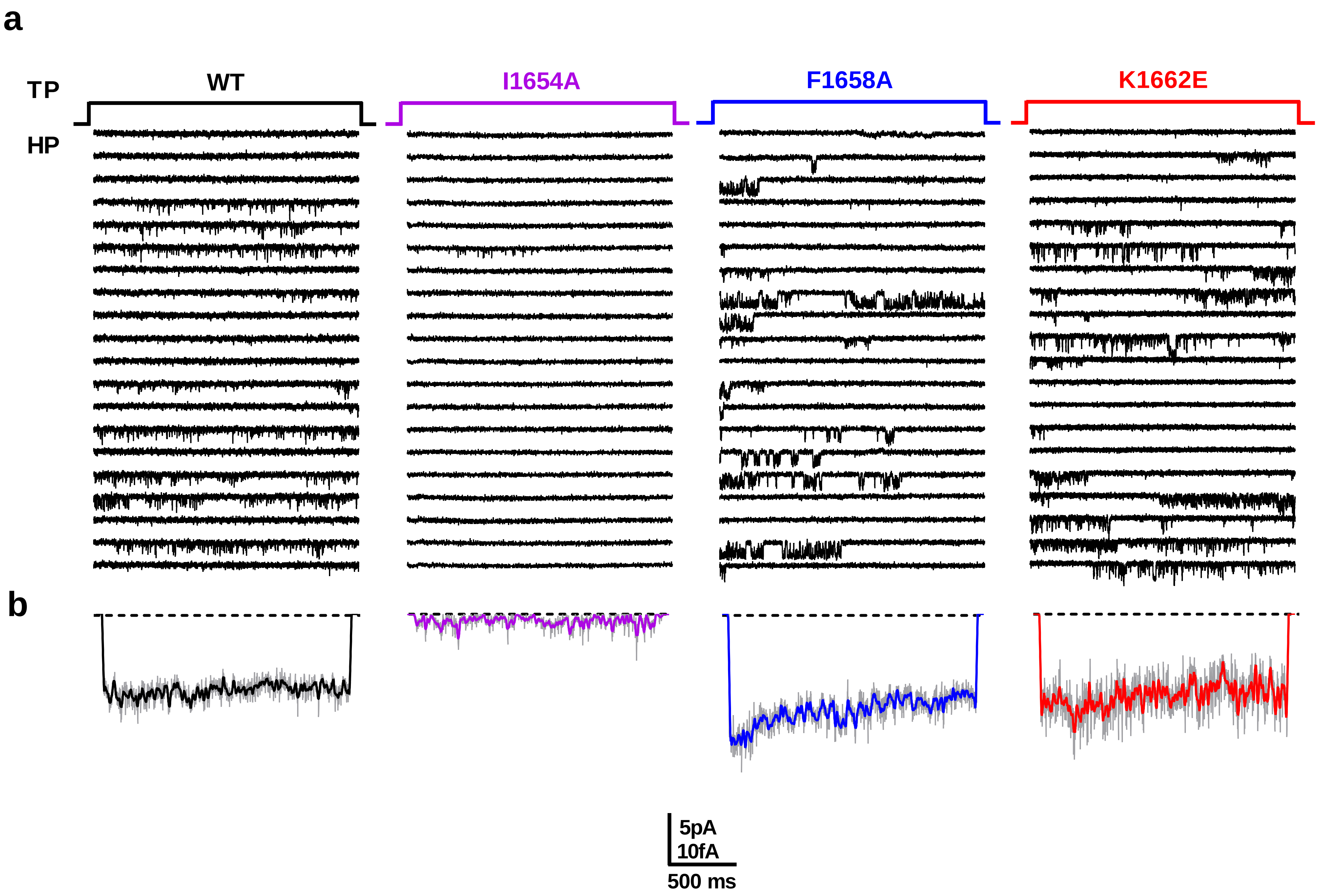

### Supplemental Figure 6

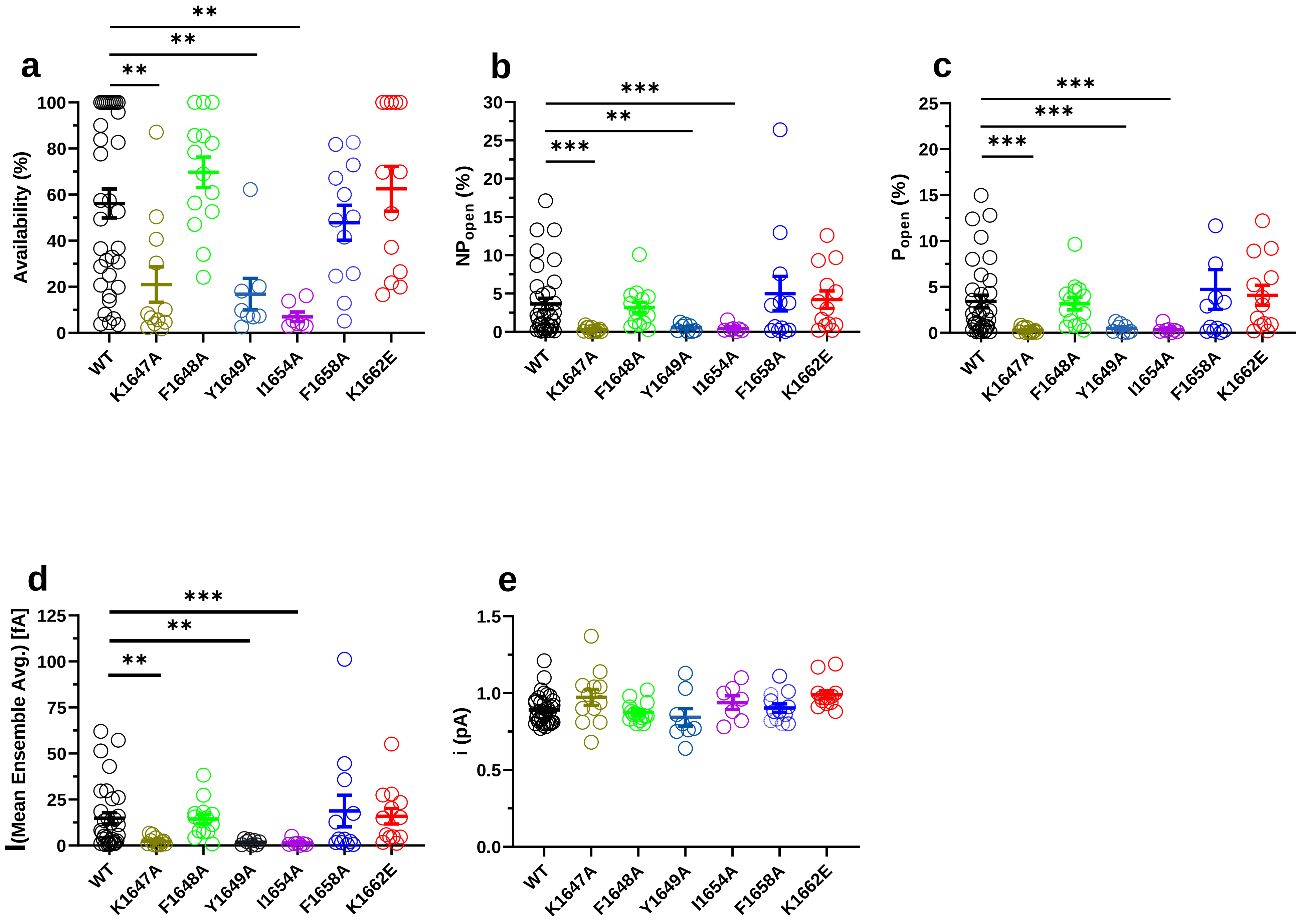

### Supplemental Figure 7

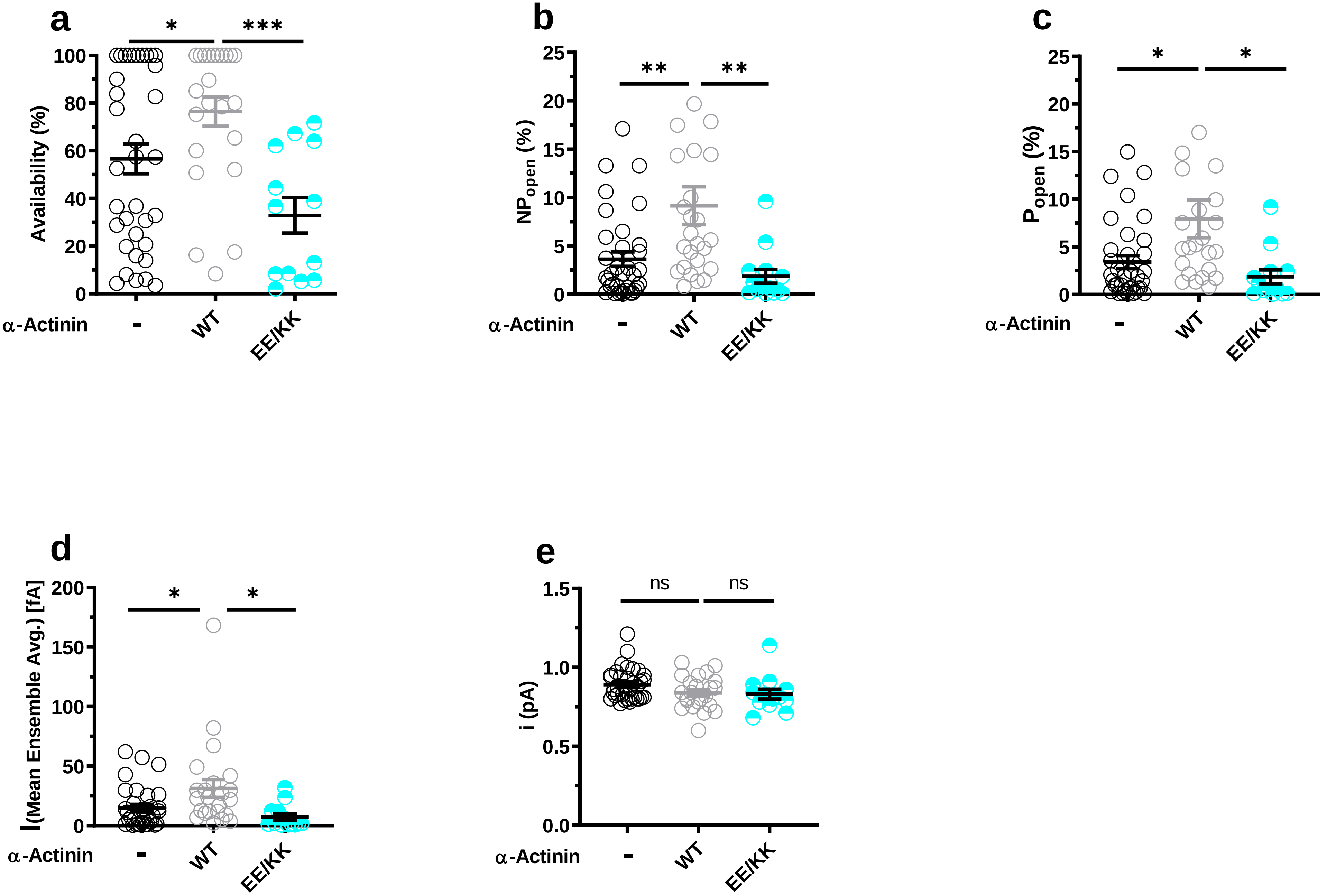

### Supplemental Figure 8

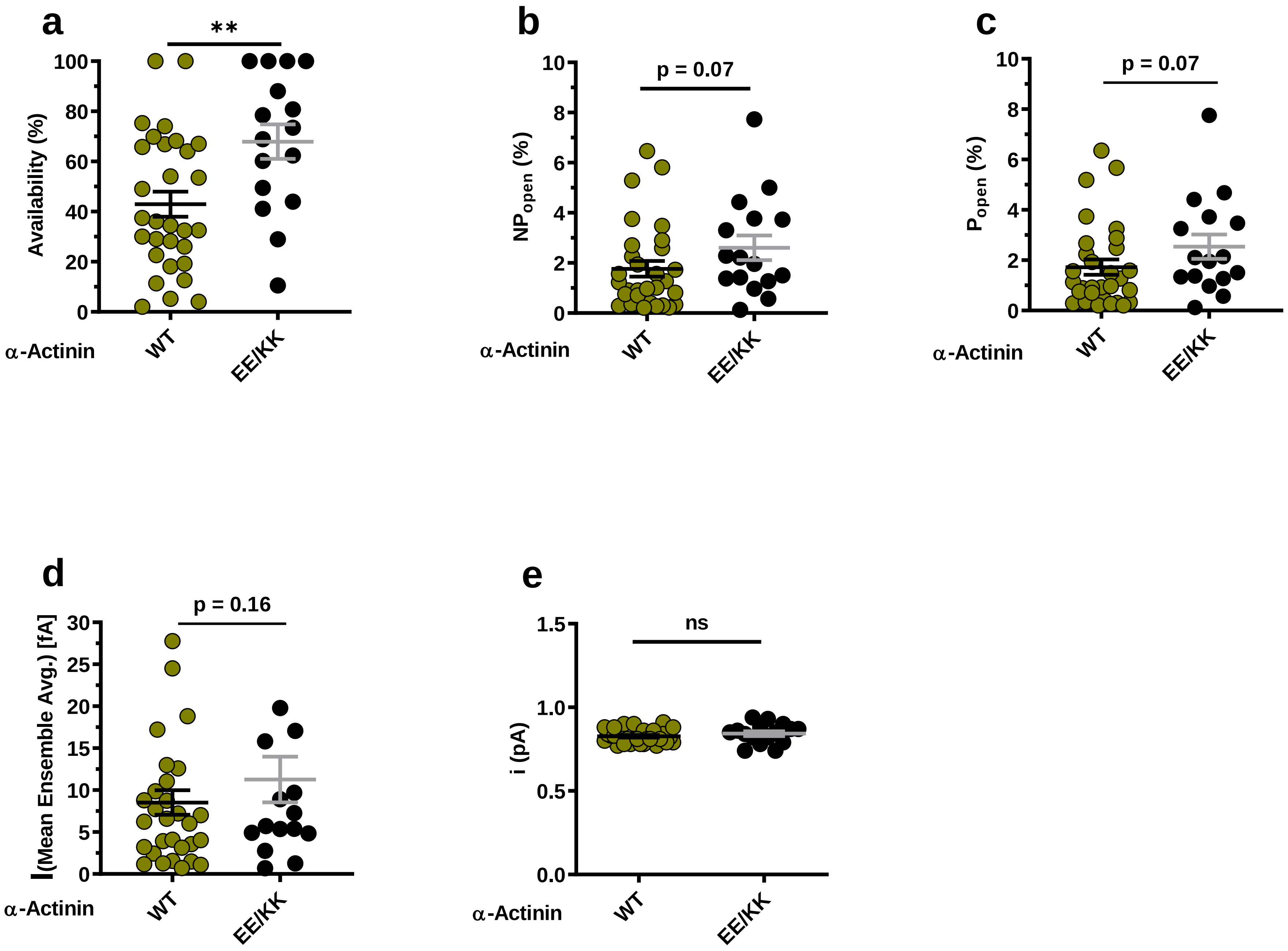

### Supplemental Figure 9

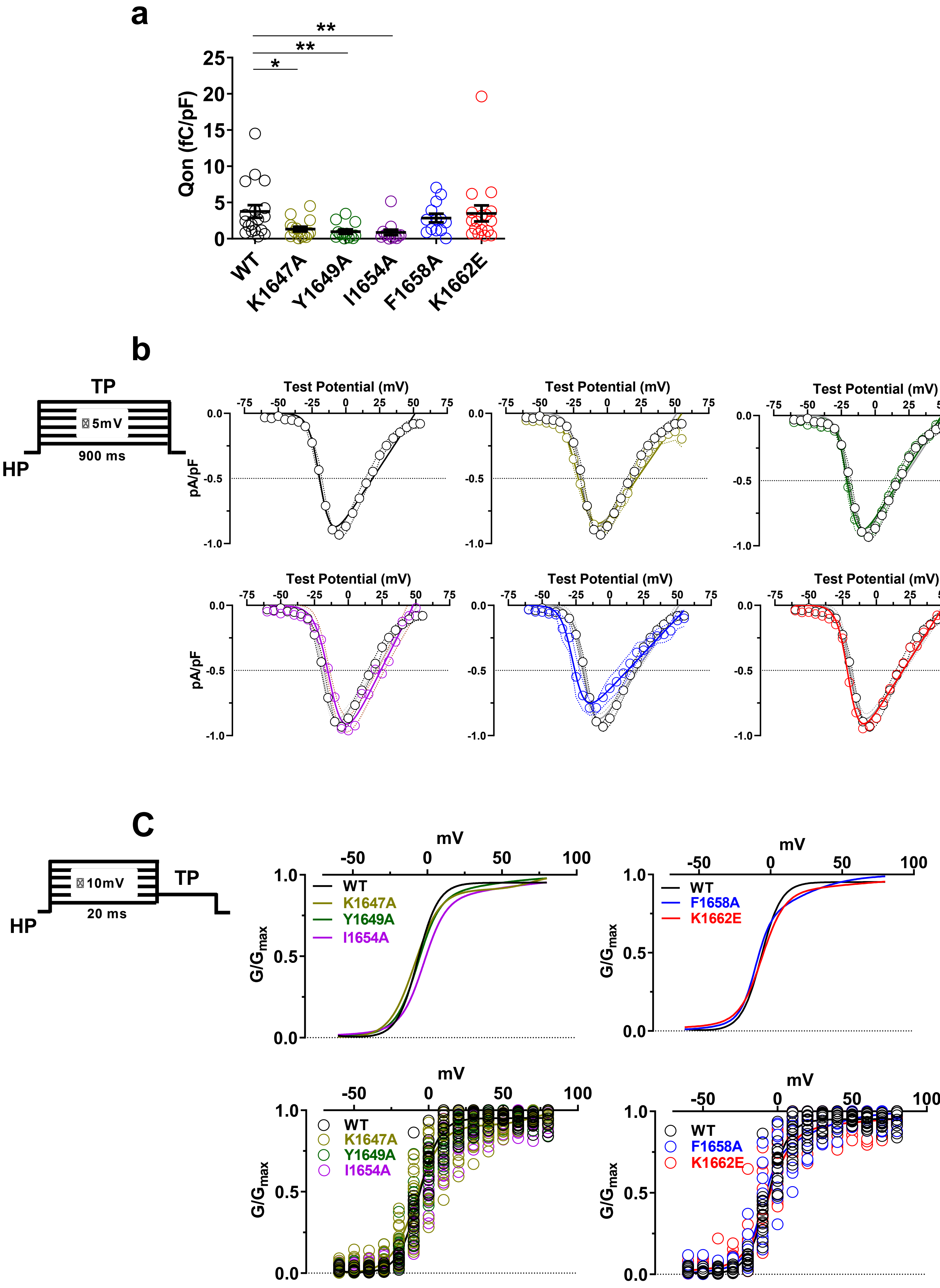

### Supplemental Figure 10

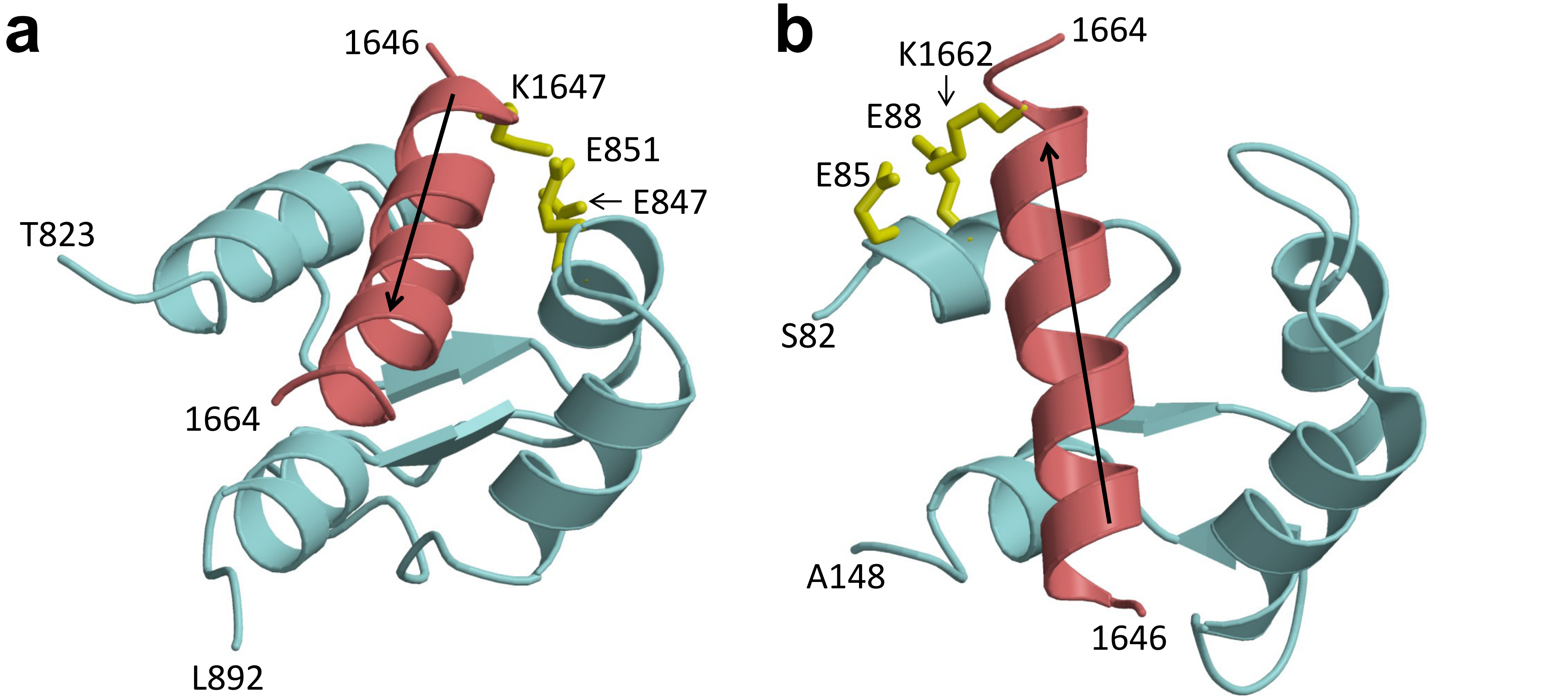


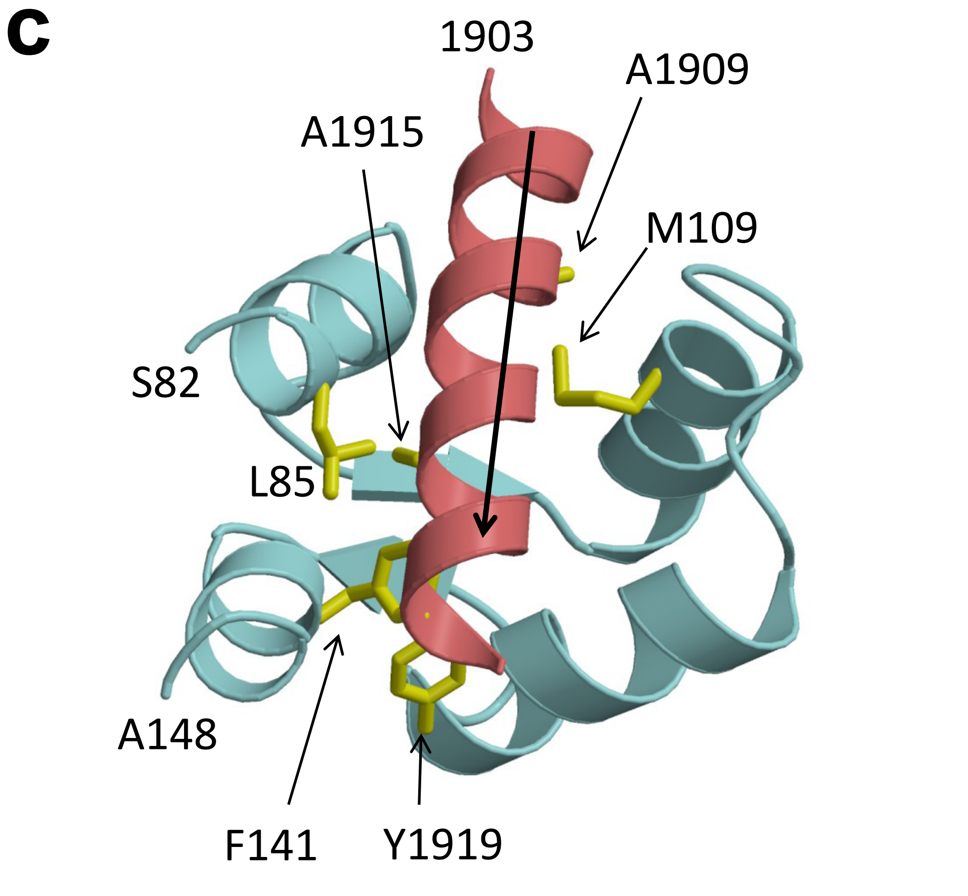


**d**

**$Na_V$1.2(1905-1922):** EEVSAIV**IQ**RAY**R**RYLLK

**$Na_V$1.5(1901-1918):** EEVSAIV**IQ**RAF**R**RHLLQ

**$Ca_V$1.2(1647-1664):** **K**FYATFL**IQ**EYF**R**KF**K**KR

### Supplemental Figure 11

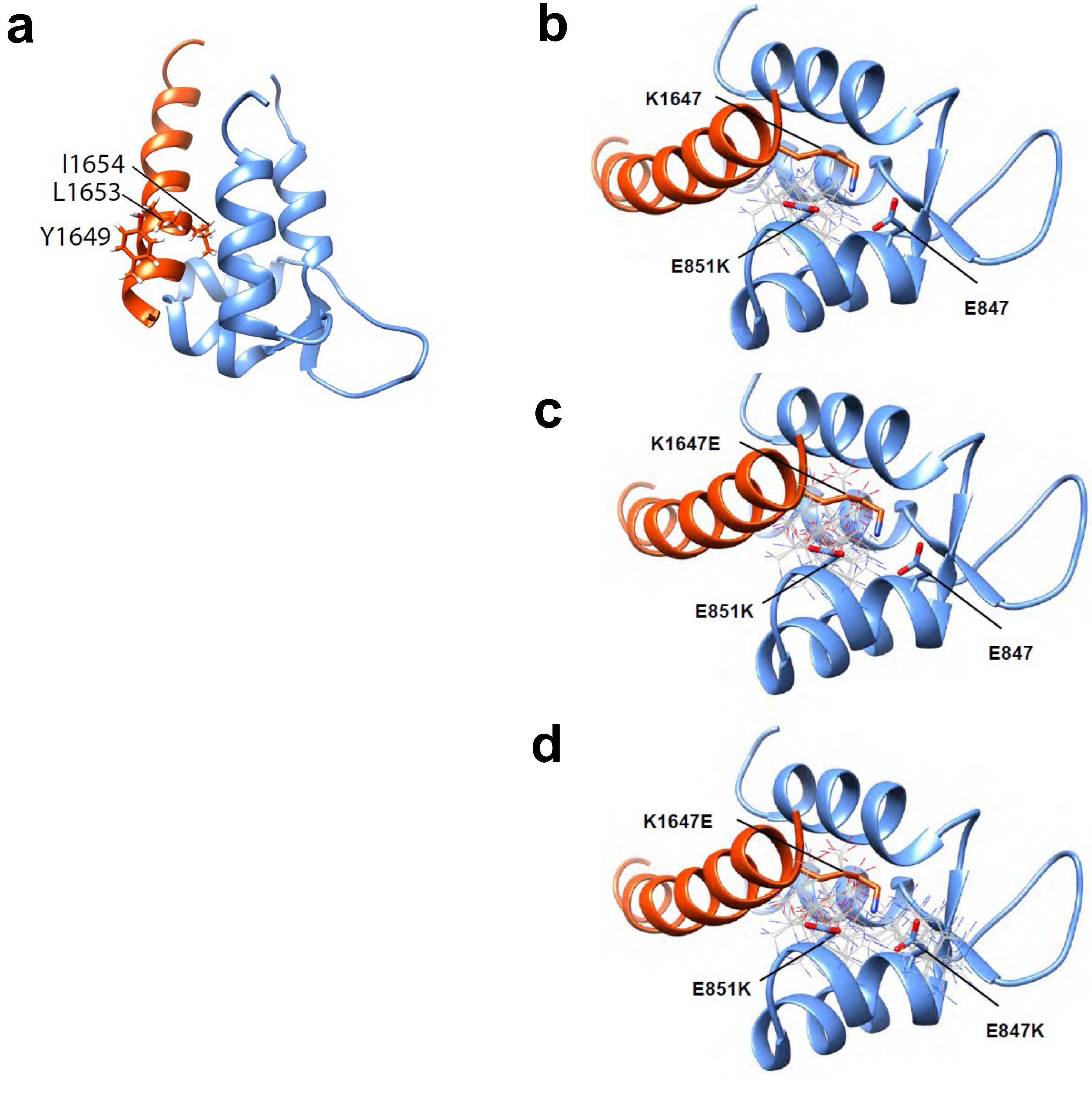
